# Dopamine receptor D2 confers colonization resistance via gut microbial metabolites

**DOI:** 10.1101/2023.03.14.532647

**Authors:** Samantha A. Scott, Jingjing Fu, Pamela V. Chang

## Abstract

The gut microbiome plays major roles in modulating host physiology. One such function is colonization resistance, or the ability of the microbial collective to protect the host against enteric pathogens^1–3^, including enterohemorrhagic *Escherichia coli* (EHEC) serotype O157:H7, an attaching and effacing (AE) food-borne pathogen that causes severe gastroenteritis, enterocolitis, bloody diarrhea, and acute renal failure (hemolytic uremic syndrome)^4,5^. Although gut microbes can provide colonization resistance by outcompeting some pathogens or modulating host defense provided by the gut barrier and intestinal immune cells, this phenomenon remains poorly understood. Emerging evidence suggests that small-molecule metabolites produced by the gut microbiota may mediate this process^6^. Here, we show that tryptophan (Trp)-derived metabolites produced by the gut bacteria protect the host against *Citrobacter rodentium*, a murine AE pathogen widely used as a model for EHEC infection^7,8^, by activation of the host neurotransmitter dopamine receptor D2 (DRD2) within the intestinal epithelium. We further find that these Trp metabolites act through DRD2 to decrease expression of a host actin regulatory protein involved in *C. rodentium* and EHEC attachment to the gut epithelium via formation of actin pedestals. Previously identified mechanisms of colonization resistance either directly affect the pathogen by competitive exclusion or indirectly by modulation of host defense mechanisms^9,10^, so our results delineate a noncanonical colonization resistance pathway against AE pathogens featuring an unconventional role for DRD2 outside the nervous system in controlling actin cytoskeletal organization within the gut epithelium. Our findings may inspire prophylactic and therapeutic approaches for improving gut health and treating gastrointestinal infections, which afflict millions globally.

## Main Text

A critical function of the gut microbiota is colonization resistance, or the ability of the gut microbiome to protect against gastrointestinal infection^1–3^. Of the estimated 500,000 small-molecule metabolites in the gut^11,12^, only a handful of microbially-produced molecules are known to protect the host from infection by enteric pathogens^13^, and the affected host pathways remain largely uncharacterized^14,15^. Enterohemorrhagic *Escherichia coli* (EHEC) serotype O157:H7 is a food-borne pathogen that causes gastroenteritis, enterocolitis, bloody diarrhea, and, in certain individuals, acute renal failure (hemolytic uremic syndrome)^4,5^. EHEC infects the host by forming attaching and effacing (AE) lesions known as actin pedestals on the gut epithelium, from which it secretes virulence factors, including Shiga toxin, into the host^16–18^. *Citrobacter rodentium*, a murine AE pathogen, is a widely used model for EHEC infection due to a highly similar infection strategy, encoded by the locus of enterocyte effacement pathogenicity island shared with EHEC and enteropathogenic *E. coli*^7,8,19^. The gut microbiota provide colonization resistance to *C. rodentium* by enhancing host defense pathways^10^ and by competing with the pathogen for nutrients within the gut^9^. Additionally, specific small-molecule metabolites produced by the gut microbiota can mediate certain colonization resistance pathways^14,15^. We were thus motivated to examine the roles of bacterial small-molecule metabolites derived from the essential amino acid tryptophan (Trp) in protecting against infection with AE pathogens because of their functions in modulating immunity^20–23^ and improving gut barrier function during inflammation^24^.

We pre-treated wildtype C57Bl/6 mice with or without a broad-spectrum antibiotic cocktail to deplete the gut microbiota, followed by a Trp-supplemented diet for 7 d, which was well-tolerated (Extended Data Fig. 1a), and infection with *C. rodentium* (Fig. 1a). Compared to mice fed conventional diet, Trp-fed mice exhibited lower colonic *C. rodentium* burden (Fig. 1b–c) and decreased colonic inflammation (Fig. 1d–f). These effects were abrogated by antibiotic treatment, suggesting that the microbiota provide protection against severe disease during infection in the presence of Trp.

**Fig. 1.**
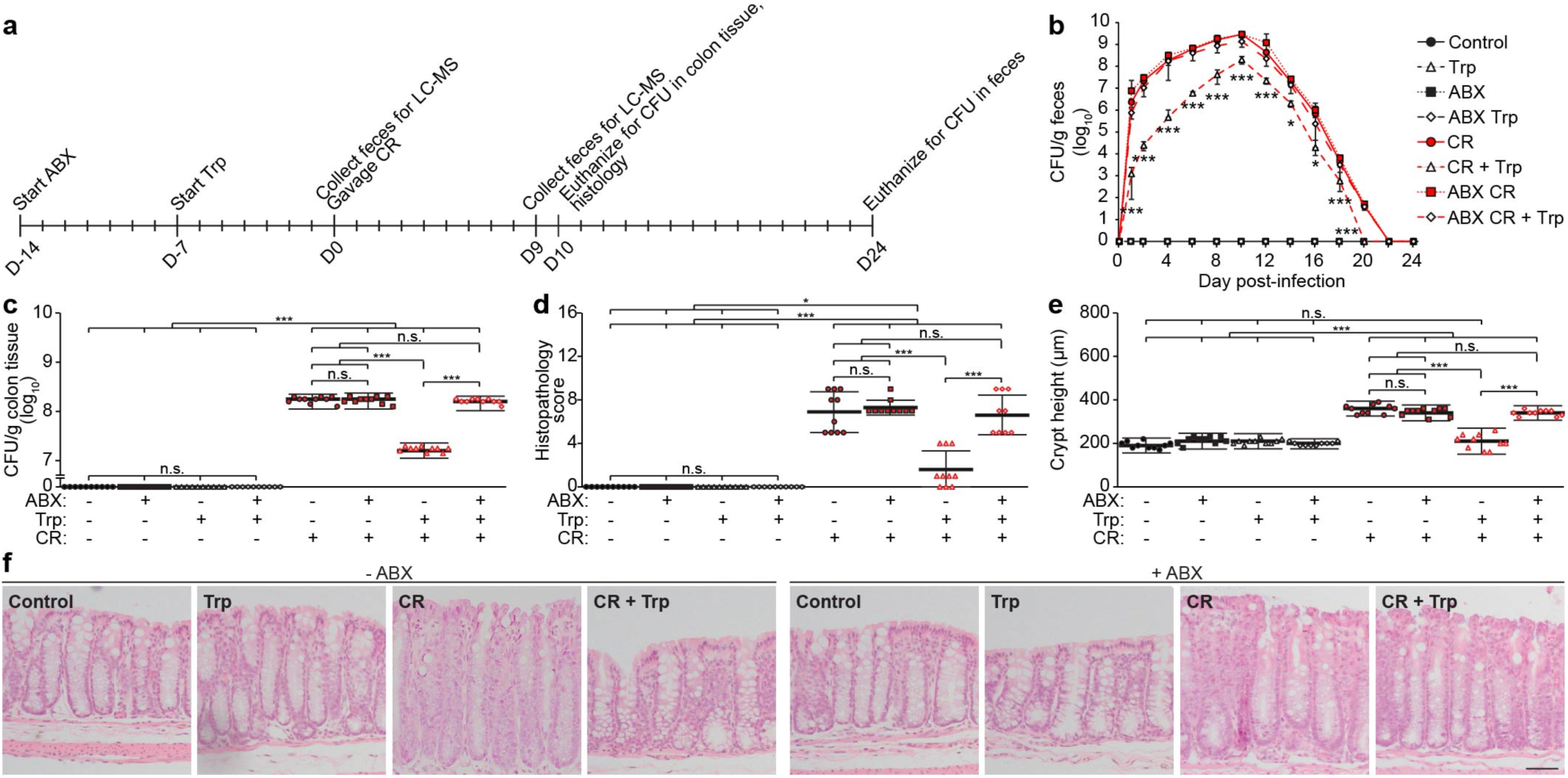
Dietary tryptophan (Trp) protects against a mouse model of EHEC infection using *Citrobacter rodentium*, strain DBS100. **a**, C57Bl/6 mice were pre-treated with antibiotics (-/+ ABX, 9 mg/kg each of metronidazole, ampicillin, and neomycin, 4.5 mg/kg of vancomycin) for 7 d, followed by conventional (2 g Trp/kg diet, ad libitum) or Trp (42 g Trp/kg diet, ad libitum) diet for 7 d. The mice were then administered *C. rodentium* (CR, oral gavage, 10^8^ colony-forming units, CFU) with continued ABX (except neomycin) and Trp feeding. Feces were collected before infection and 9 d post-infection to determine Trp metabolite levels (see Extended Data Fig. 1a, b). **b, c**, Bacterial load in (**b**) feces and (**c**) colon tissue was measured (**b**) at 1–2 d intervals for 24 d post-infection and (**c**) at the peak of infection, 10 d post-infection. **d–f**, Colon sections were stained with H&E and (**d**) blindly scored for submucosal edema (0-3), goblet cell depletion (0-3), epithelial hyperplasia (0-3), epithelial integrity (0-4), and neutrophil and mononuclear cell infiltration (0-3). Data are expressed as the sum of these individual scores (0-16). See Methods for full description of scoring rubric. (**e**) Crypt heights were measured. (**f**) Representative images. Scale bar: 50 μm. Data are representative of at least 3 independent experiments, n=10 mice per group, bars = mean, error bars = standard deviation. One-way ANOVA followed by post-hoc Tukey’s test: *p<0.05, ***p<0.001, n.s. = not significant.

To identify specific Trp metabolites that mediate the effects of the Trp diet, we performed targeted mass spectrometry (MS)-based metabolomics before and after *C. rodentium* infection during Trp feeding and found that the most highly abundant metabolites were indole, indole-3-ethanol (IEt), indole-3-pyruvate (IPyA), and indole-3-aldehyde (I3A) (Extended Data Fig. 1b). Though indole can decrease EHEC and *C. rodentium* virulence, the effects and mechanisms of action of IEt, IPyA, and I3A on these pathogens remain mostly unknown^25–27^.

To determine the roles of these Trp metabolites on *C. rodentium* infection, we pre-treated C57Bl/6 mice with or without antibiotics to prevent microbial metabolism of the individual Trp metabolites, followed by administration of IEt, IPyA, or I3A for 2 d prior to and during infection (Fig. 2a). Treatment with the individual metabolites lowered colonic *C. rodentium* burden (Fig. 2b–c) and decreased colonic inflammation (Fig. 2d–f). We verified that levels of IEt, IPyA, and I3A in the colon were increased following metabolite administration and did not change with antibiotic treatment (Extended Data Fig. 1c–e). We also determined that Trp diet, IEt, IPyA, and I3A did not significantly alter microbial composition by 16S rRNA gene analysis (Extended Data Fig. 2). Thus, these microbial Trp metabolites mediate colonization resistance against *C. rodentium*.

**Fig. 2.**
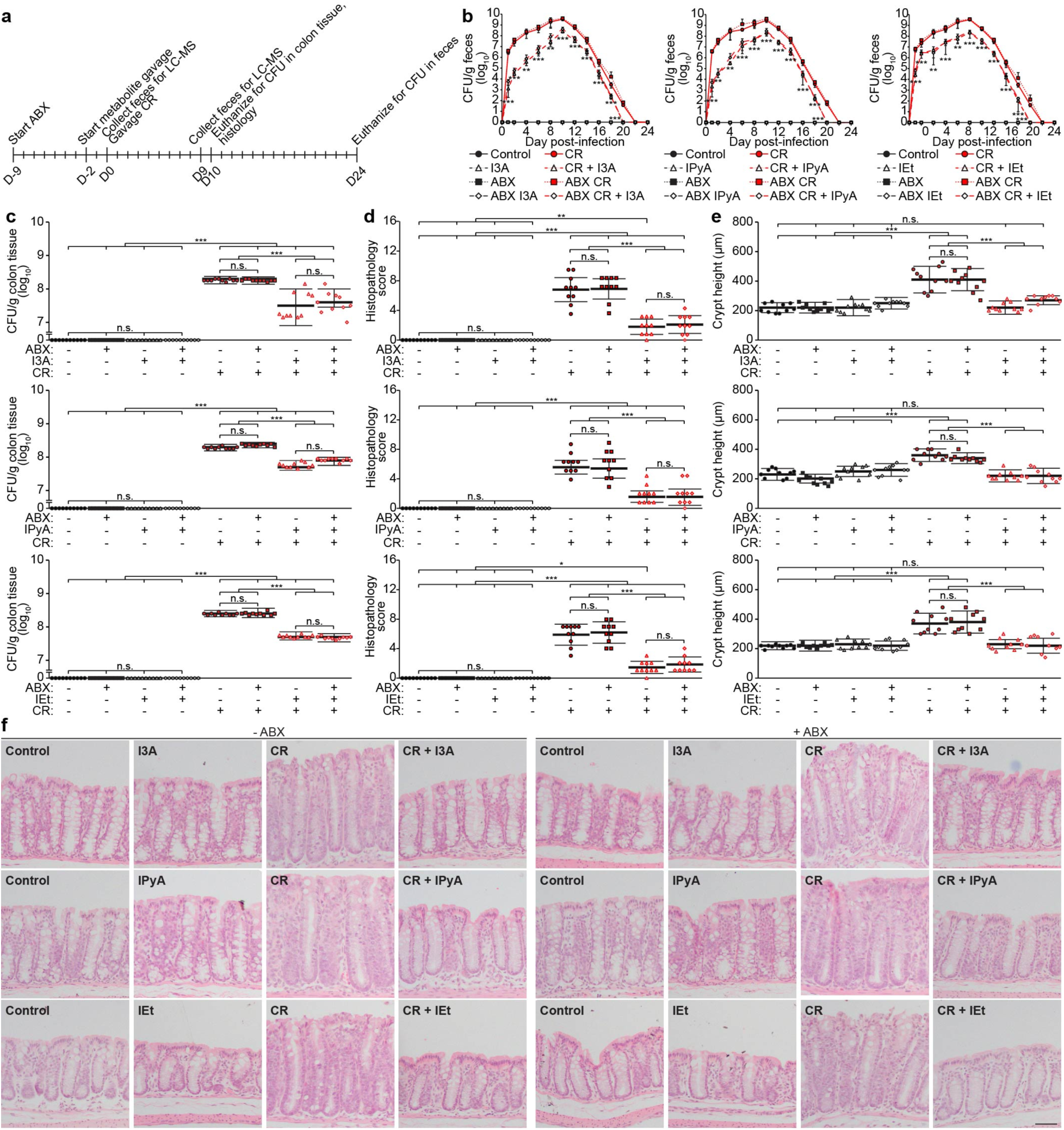
The Trp metabolites I3A, IPyA, and IEt protect against *Citrobacter rodentium* infection in mice. **a**, C57Bl/6 mice were pre-treated with antibiotics (ABX) for 7 d, followed by Trp metabolites, I3A (1000 mg/kg), IPyA (2900 mg/kg), or IEt (600 mg/kg) by oral gavage daily for 2 d. The mice were then administered *C. rodentium* (CR, oral gavage, 10^8^ CFU) with continued ABX (except neomycin) and metabolite treatment. Feces were collected before infection and 9 d post-infection to determine Trp metabolite levels (see Extended Data Fig. 1c–e). **b, c**, Bacterial load in (**b**) feces and (**c**) colon tissue was measured (**b**) at 1–2 d intervals for 24 d post-infection and (**c**) at the peak of infection, 10 d post-infection. **d–f**, Colon sections were stained with H&E and (**d**) blindly scored as described in Fig. 1. (**e**) Crypt heights were measured. (**f**) Representative images. Scale bar: 50 μm. Data are representative of at least 3 independent experiments, n=10 mice per group, bars = mean, error bars = standard deviation. One-way ANOVA followed by post-hoc Tukey’s test: *p<0.05, **p<0.01, ***p<0.001, n.s. = not significant.

To determine if IEt, IPyA, and I3A affect the pathogens directly, we treated *C. rodentium* with each metabolite and found that they did not affect bacterial growth and virulence factor expression (Extended Data Fig. 3a–e). We also treated EHEC with IEt, IPyA, and I3A and found that each metabolite did not affect this pathogen as well (Extended Data Fig. 3f–k). Because the metabolites did not directly affect bacterial growth or virulence in vitro, we reasoned that IEt, IPyA, and I3A may mediate colonization resistance primarily via host pathways.

To identify potential host receptors for these metabolites, we utilized Similarity Ensemble Approach^28^, a computational tool for predicting protein targets of small molecules, with IEt, IPyA, and I3A as ligands. All three metabolites were predicted to bind the dopamine receptors D2 (DRD2), D3 (DRD3), and D4 (DRD4), G protein-coupled receptors (GPCRs) with canonical roles as neurotransmitter receptors in the central and peripheral nervous systems^29,30^. We found that DRD2–4 inhibition using the pan-inhibitor haloperidol^31^ reduced the effects of the metabolites in decreasing bacterial attachment via actin pedestals during EHEC infection of a human intestinal epithelial cell (IEC) line, Caco-2 (Extended Data Fig. 4a, b). Further, we found that this effect depended exclusively on *Drd2*, using CRISPR/Cas9 knockout of *Drd2*, *Drd3*, and *Drd4* in Caco-2 cells during EHEC infection (Extended Data Fig. 4c–f). Critically, we found that IEt, IPyA, and I3A are bona fide DRD2 ligands, with potencies similar to the endogenous ligand, dopamine, using both the GloSensor and Tango GPCR assays, which quantify cyclic AMP and arrestin recruitment, respectively (Extended Data Fig. 5)^32,33^. We also pre-treated C57Bl/6 mice with or without the antibiotic cocktail, followed by dopamine and *C. rodentium* infection. We found that dopamine had similar effects, albeit with lower potency likely due to decreased bioavailability^34^, compared to the Trp diet, IEt, IPyA, and I3A in decreasing morbidity during infection (Extended Data Fig. 6a–f). Further, treatment of mice with a DRD2 agonist, sumanirole, had similar effects to the Trp metabolites (Extended Data Fig. 6j–n), whereas a DRD2 antagonist, L-741,626, blocked the effects of the Trp diet (Extended Data Fig. 6r–v). Collectively, these data suggest that DRD2 may be a host receptor in the gut that recognizes the Trp metabolites.

We first assessed whether DRD2 expression in the gut epithelium, which expresses this receptor and is the initial target tissue of AE pathogens, mediates the effects of the Trp metabolites in promoting colonization resistance to *C. rodentium* infection. We generated *Drd2*fl/fl x Villin (Vil)-Cre mice, which lack *Drd2* expression in IECs (Extended Data Fig. 7a), and treated these mice with Trp diet, IEt, IPyA, or I3A, followed by infection with *C. rodentium* (Fig. 3a, Extended Data Figs. 7b–c and 8). We found that *Drd2*fl/fl control littermates that were treated with Trp diet or individual metabolites exhibited decreased colonic *C. rodentium* burden (Figs. 3b–c and Extended Data Fig. 9a–b), colonic inflammation as measured by histology (Figs. 3d–f and Extended Data Fig. 9c–e), and immune cell infiltration (Extended Data Figs. 10–13) and inflammatory mediator production (Extended Data Fig. 14). However, these effects were abrogated in the *Drd2*fl/fl x Vil-Cre mice, indicating that DRD2 expression in IECs mediates the effects of the Trp metabolites. Because serotonin can affect neuronal dopaminergic signaling^35,36^ and gut luminal serotonin decreases *C. rodentium* virulence ^37^, we treated mice with inhibitors of serotonin selective reuptake transporter or tryptophan hydroxylase to elevate or decrease serotonin levels, respectively, and found that the Trp diet mediated colonization resistance to *C. rodentium* irrespective of serotonin levels. (Extended Data Fig. 15).

**Fig. 3.**
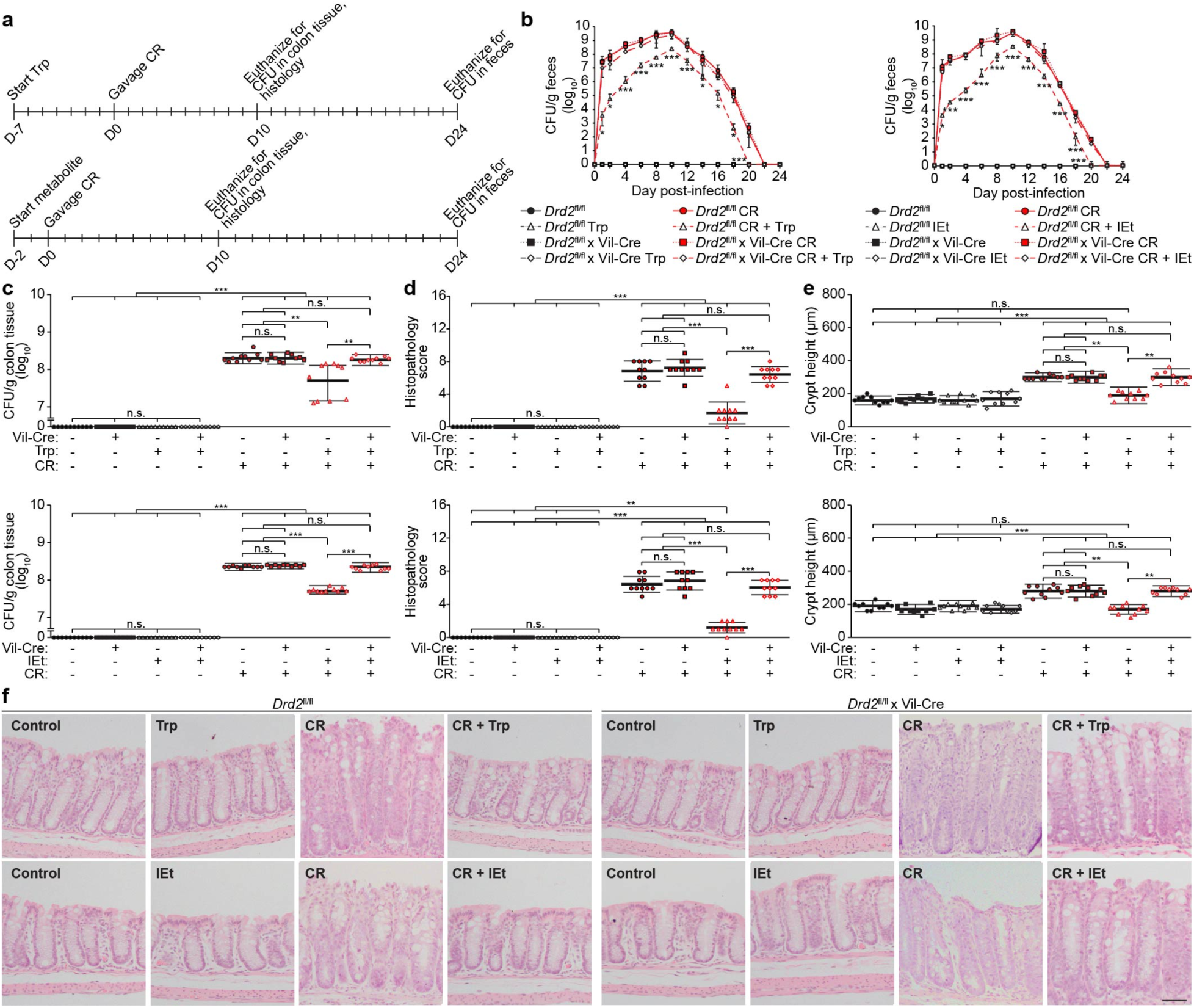
Effects of the Trp diet and metabolites in protecting against *Citrobacter rodentium* infection depend on dopamine receptor D2 (DRD2) in intestinal epithelial cells (IECs). **a**, *Drd2*fl/fl x Villin (Vil)-Cre or *Drd2*fl/fl mice were fed a conventional (2 g Trp/kg diet, ad libitum) or Trp (42 g Trp/kg diet, ad libitum) diet for 7 d or Trp metabolites (shown: IEt (600 mg/kg); see Extended Data Fig. 9 for I3A and IPyA), by oral gavage daily for 2 d, and then infected with *C. rodentium* (CR, oral gavage, 10^8^ CFU) with continued Trp feeding or metabolite treatment. **b, c**, Bacterial load in (**b**) feces and (**c**) colon tissue was measured (**b**) every 1–2 d for 24 d post-infection and (**c**) at the peak of infection, 10 d post-infection. **d–f**, Colon sections were stained with H&E and (**d**) blindly scored as described in Fig. 1. (**e**) Crypt heights were measured. (**f**) Representative images. Scale bar: 50 μm. Data are representative of at least 3 independent experiments, n=10 mice per group, bars = mean, error bars = standard deviation. One-way ANOVA followed by post-hoc Tukey’s test: *p<0.05, **p<0.01, ***p<0.001, n.s. = not significant.

We further examined the effects of the Trp diet, IEt, IPyA, and I3A in *C. rodentium*-infected *Drd2*fl/fl mice lacking *Drd2* expression in immune cells that also express DRD2 and are involved in host immunity against this pathogen, including macrophages (MPs), dendritic cells (DCs), and CD4^+^ T cells, using *Drd2*fl/fl x LysM-Cre, *Drd2*fl/fl x CD11c-Cre, and *Drd2*fl/fl x CD4-Cre mice, respectively. The effects of the Trp diet were not eliminated in the mice lacking *Drd2* in MPs, DCs, and CD4^+^ T cells, indicating that the metabolites act via DRD2 expressed in IECs (Extended Data Fig. 16).

We then sought to determine mechanisms by which DRD2 mediates colonization resistance in IECs. Because the actin regulatory protein N-WASP constitutes an initial host protein hijacked by AE pathogens during intestinal colonization that is required for pedestal formation and mucosal colonization of *C. rodentium*^17,18^, we examined the effects of the Trp diet and metabolites on the formation of actin pedestals by this protein during infection. We found that the Trp diet, IEt, IPyA, and I3A all reduced pedestal formation during *C. rodentium* infection in wildtype or *Drd2*fl/fl mice and that this decrease was eliminated in antibiotic-treated (Fig. 4a and Extended Data Fig. 9g) or *Drd2*fl/fl x Vil-Cre mice (Fig. 4b and Extended Data Fig. 9f). We further found that the Trp diet and metabolites decreased N-WASP protein levels in the gut epithelium before and during *C. rodentium* infection in *Drd2*fl/fl mice, and these effects were also abrogated in the *Drd2*fl/fl x Vil-Cre mice (Extended Data Fig. 9h–i). Dopamine and the DRD2 agonist sumanirole had similar effects as the Trp diet and metabolites on actin pedestal formation and N-WASP levels in *C. rodentium* infection (Extended Data Fig. 6g-i, o–q), whereas the DRD2 antagonist L-741,626 inhibited the effects of the Trp diet (Extended Data Fig. 6w–y). Further, we determined that IEt, IPyA, and I3A act via DRD2 to protect against EHEC infection of Caco-2 monolayers by a similar mechanism (Fig. 4c, w–x and Extended Data Fig. 4f, k–l), suggesting that our findings may be generalizable to related AE human pathogens.

**Fig. 4.**
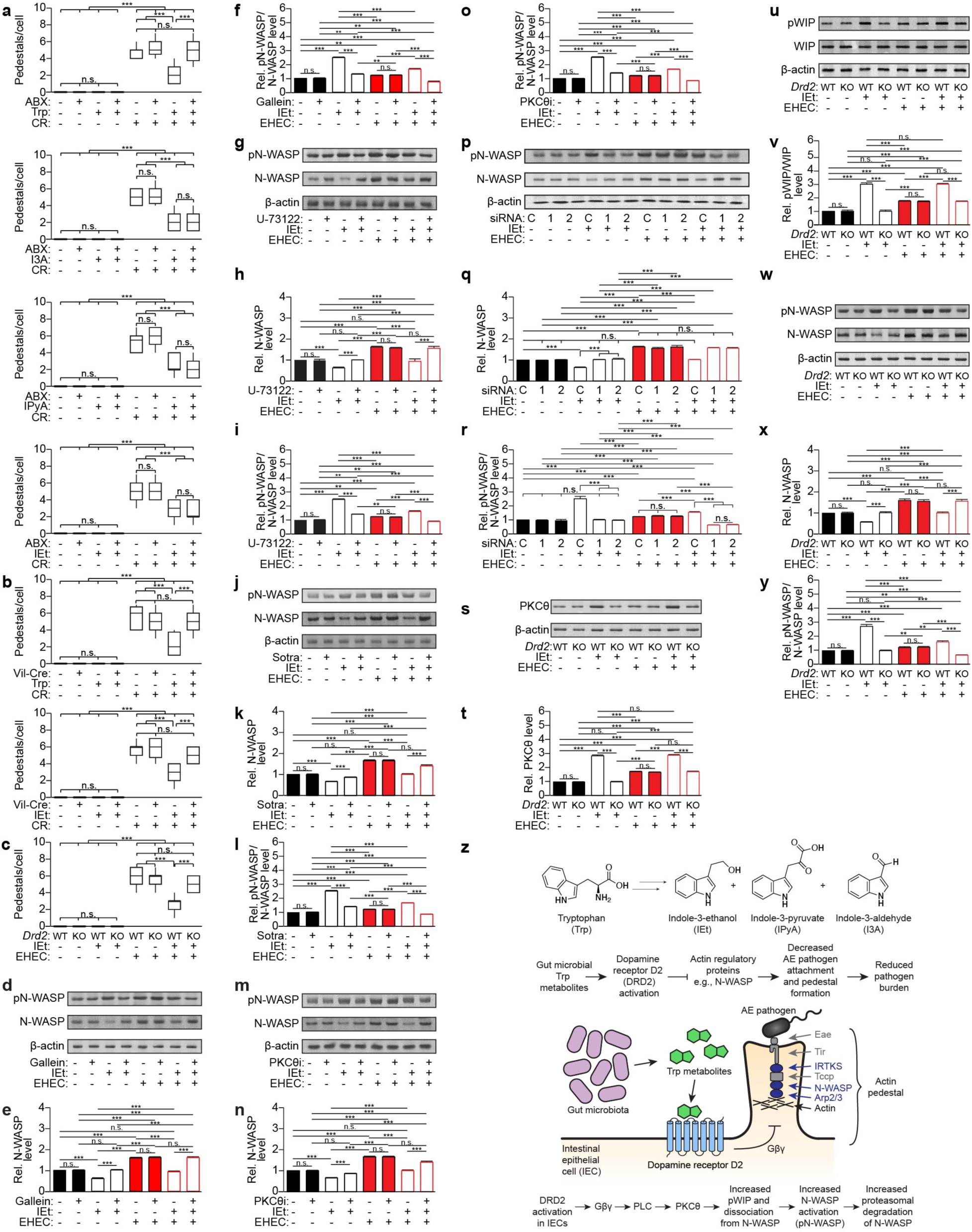
Tryptophan (Trp) metabolites decrease actin pedestal formation via DRD2 in IECs during *Citrobacter rodentium* and EHEC O157:H7 infection. **a**, C57Bl/6 mice were pre-treated with antibiotics (-/+ ABX, 9 mg/kg each of metronidazole, ampicillin, and neomycin, 4.5 mg/kg of vancomycin) for 7 d, followed by conventional (2 g Trp/kg diet, ad libitum) or Trp (42 g Trp/kg diet, ad libitum) diet for 7 d or Trp metabolites, I3A (1000 mg/kg), IPyA (2900 mg/kg), or IEt (600 mg/kg) by oral gavage daily for 2 d. The mice were then administered *C. rodentium* (CR, oral gavage, 10^8^ CFU) with continued ABX (except neomycin) and Trp feeding or metabolite treatment. **b**, *Drd2*fl/fl x Villin (Vil)-Cre or *Drd2*fl/fl mice were fed a conventional (2 g Trp/kg diet, ad libitum) or Trp (42 g Trp/kg diet, ad libitum) diet for 7 d or Trp metabolites, I3A (1000 mg/kg), IPyA, (2900 mg/kg), or IEt (600 mg/kg) by oral gavage daily for 2 d, and then infected with *C. rodentium* (CR, oral gavage, 10^8^ CFU) with continued Trp feeding or metabolite treatment (shown: IEt). **c**, Caco-2 monolayers (*Drd2* WT vs. KO) were pre-treated with metabolites (100 μM) for 2 d and then infected with EHEC O157:H7 (MOI 50) for 16 h. (**a–c**) Samples were stained with DAPI and Alexa Fluor 647-phalloidin. Pedestal formation = # of pedestals per host or Caco-2 cells (ABX + Trp diet, n = 146–177; ABX + I3A, 121–184; ABX + IPyA, 118–151; ABX + IEt, 118–175; Vil-Cre: Trp diet, n=110–186; IEt, n=100–130, *Drd2* WT vs KO Caco-2 cells: IEt, n = 130–188). (**d–o**) Caco-2 cells were pre-treated with the indicated inhibitors (Gβγ inhibitor gallein, 10 μM; PLC inhibitor U-73122, 10 μM; pan-PKC inhibitor sotrastaurin (Sotra), 5 μM; or PKC-8 inhibitor (PKC8i), 5 μM), followed by metabolites (100 μM) for 24 h and then infected with EHEC O157:H7 (MOI 50) for 12 h. (**p–r**) PKC-8 knockdown was performed using two different siRNA duplexes (1 and 2) or negative control siRNA (C). (s–y) Caco-2 cells (*Drd2* WT vs KO) were pre-treated with metabolites (100 μM) for 24 h and then infected with EHEC O157:H7 (MOI 50) for 12 h. (**d–y**) Cell lysates were analyzed by Western blotting with the indicated antibodies. (**e, f, h, i, k, l, n, o, q, r, t, v, x, y**) Densitometry was performed using FIJI, n = 3. z, Model of gut microbial Trp metabolite activation of DRD2 in IECs and reduced pathogen burden. Blue = host actin regulatory proteins. Data are representative of at least 3 independent experiments, n=10 mice per group, bars = mean, error bars = standard deviation. One-way ANOVA followed by post-hoc Tukey’s test: **p<0.01, ***p<0.001, n.s. = not significant.

To determine the molecular mechanism underlying how DRD2 activation in IECs leads to decreased pedestal formation and AE pathogen attachment, we assessed the dependence of the metabolite-induced effects on signaling pathways downstream of DRD2 using inhibitors of Gαi, Gβγ, and β-arrestin signaling. We found that inhibition of Gβγ, but not Gαi or β-arrestin signaling (Extended Data Fig. 17), prevented the metabolite-induced decrease in N-WASP levels and increase in N-WASP phosphorylation, the latter of which leads to its subsequent proteasomal degradation^38^ (Fig. 4d–f and Extended Data Fig. 18a, f–h). Blockade of phospholipase C (PLC) and protein kinase C (PKC)-8, which are downstream of Gβγ^30^, also eliminated these effects of the Trp metabolites on N-WASP, including prevention of the phosphorylation of Wiskott-Aldrich syndrome protein (WASP) interacting protein (WIP), a PKC-8 substrate^39^ whose phosphorylation causes N-WASP^40^ activation, leading to subsequent proteasomal degradation of N-WASP^38^ (Fig. 4g–r, Extended Data Fig. 18b–e, i–p, and Extended Data Fig. 19–21). Critically, this pathway was dependent on DRD2, indicating that the activation of DRD2 in IECs by Trp metabolites leads to decreased N-WASP levels, and hence decreased AE pathogen attachment and colonization, via sequential activation of Gβγ, PLC, and PKC-8 (Fig. 4s–y, Extended Data Fig. 4g–m).

We demonstrate that metabolites produced by the gut microbiota mediate colonization resistance against AE pathogens, including EHEC and *C. rodentium*, by decreasing the expression of a key host actin regulatory protein hijacked by these pathogens during initial colonization (Fig. 4z). Critically, these Trp metabolites activate the neurotransmitter receptor DRD2 expressed on IECs to control the expression of these host proteins. These findings represent a unique pathway of colonization resistance that differs from previously described immune mediated pathways^41,42^. Interestingly, a recent high-throughput screen showed that DRD2 can be activated by additional microbial metabolites^43^, suggesting that this receptor may function more generally as an aromatic amino acid metabolite sensor and an important mediator of host-microbe interactions in the gut. Our studies reveal that, beyond its canonical roles in the nervous system, DRD2 plays an unconventional role in conferring host defense in the gut. DRD2 and its downstream pathways could serve as targets for prophylactics or therapeutics for improving gut health and treating gastrointestinal infection with certain AE pathogens.

## Methods

### Bacterial strains and culture

EHEC EDL-933 and TUV93-0 was a gift of Tobi Doerr (Cornell University) and John Leong (Tufts University), respectively. *C. rodentium* DBS100 was obtained from Gregory Sonnenberg (Weill Cornell Medicine). Difco MacConkey agar was purchased from BD Biosciences. Bacteriological agar was purchased from VWR, yeast extract and tryptone were purchased from IBI Scientific, and sodium chloride was purchased from Fisher Scientific.

### Tissue culture

Caco-2 and HEK 293T cells were obtained from the American Tissue Culture Company and cultured according to their guidelines. HEK 293T cells stably expressing a GloSensor cAMP reporter were obtained from Thomas Gardella (Harvard Medical School). DMEM, RPMI-1640, penicillin/streptomycin (P/S), 0.05% trypsin, L-glutamine, HEPES, and DPBS were obtained from Corning, and Seradigm Premium Grade Fetal Bovine Serum (FBS) and Transfectagro were purchased from VWR. Sodium pyruvate was obtained from Lonza, and 2-mercaptoethanol was purchased from Sigma-Aldrich. Cell culture transwell inserts (transparent PET membrane, 12-well, 0.4 µm pore size) and Falcon 12-well companion plates were obtained from BD Falcon. Lipofectamine 2000, lipofectamine RNAiMAX, and DMEM (low glucose) were purchased from Thermo Fisher Scientific. Polybrene and puromycin were purchased from EMD Millipore Sigma.

### Metabolites and compounds

L-Tryptophan (Trp) was purchased from Chem-Impex International, Inc. Indole-3-aldehyde (I3A) and indole-3-pyruvate (IPyA) were obtained from Biosynth. Tryptophol (IEt), indole-3-propionate (IPA), and dopamine hydrochloride (DA) were obtained from Alfa Aesar. Kynurenine (Kyn) and sotrastaurin (Sotra) were purchased from Cayman Chemical Company. Indole (IND), indole-3-acetamide (IAM), DL-indole-3-lactate (ILA), 5-hydroxytryptamine (5HT), and indole-3-acetic acid (IAA), and gallein were purchased from Sigma Aldrich. Tryptamine hydrochloride (TrA) was obtained from TCI America, and indole-3-acrylate (IA) was purchased from Santa Cruz Biotechnology. Sumanirole maleate (SUM), L-741,626, fluoxetine hydrochloride (Prozac), and 4-chloro-DL-phenylalanine methyl ester hydrochloride (PCPA) were purchased from Neta Scientific. Pertussis toxin (PTx) was purchased from Thermo Fisher Scientific. Barbadin was obtained from MedChem Express. PKC-θ inhibitor (PKCθi) and MG-132 were purchased from Selleck Chemicals.

### Histology

Hematoxylin was purchased from VWR, and eosin Y was obtained from Acros Organics. Canada Balsam was obtained from Ward’s Science, and xylenes was obtained from Macron Fine Chemicals.

### Antibiotics

Ampicillin and vancomycin hydrochloride were purchased from Sigma Aldrich. Neomycin sulfate hydrate and metronidazole were obtained from Alfa Aesar.

### Western blotting

DC Protein Assay kit was purchased from BioRad. SuperSignal West Pico Chemiluminescent Substrate was purchased from Thermo Fisher Scientific. Bovine serum albumin (BSA) was purchased from VWR, and non-fat dry milk was purchased from Laboratory Product Sales. Protease inhibitor (cOmplete) tablets were obtained from Roche. Sodium β-glycerophosphate was obtained from Alfa Aesar, and sodium orthovanadate was purchased from MP Biomedicals. Sodium fluoride was purchased from Chem-Impex International, Inc., and sodium pyrophosphate decahydrate was purchased from Fisher Scientific. Anti-DRD2 (clone B-10, sc-5303), anti-DRD3 (clone 9F4, sc-136170), anti-DRD4 (clone 2B9, sc-136169), anti-PKC-θ (clone E-7, sc-1680), anti-WIP (clone A-7, sc-271113), and anti-N-WASP (clone C-1, sc-271484) were purchased from Santa Cruz Biotechnology. Alexa Fluor 647 mouse anti-WIP (pSer488, clone K32-824, 558674) was purchased from BD Biosciences. Anti-N-WASP (pTyr256, WP2601) was purchased from ECM Biosciences. Anti-mouse horse radish peroxidase-conjugate (170-5947) and anti-GAPDH (clone 6C5, MCA4739) were purchased from BioRad. Anti-β-actin (clone C-4, ICN691002) was purchased from Thermo Fisher Scientific. Anti-α-tubulin (clone B512, T5168) antibody was purchased from EMD Millipore Sigma.

### Immunofluorescence

Paraformaldehyde (PFA) 32% solution, EM grade, was purchased from Electron Microscopy Sciences, and DAPI Prolong Diamond was purchased from Thermo Fisher Scientific. Fisher HealthCare Tissue Plus OCT Compound was obtained from Fisher Scientific. Donkey anti-mouse Alexa Fluor 594 (A21293) and Alexa Fluor 647-phalloidin (A22287) were purchased from Thermo Fisher Scientific.

### LC-MS

LC-MS grade methanol, water, acetonitrile, and formic acid (FA) were obtained from Fisher Scientific.

### qPCR

RNABee was obtained from Tel-Test, Inc. Diethylpyrocarbonate (DEPC) and chloroform were purchased from Sigma Aldrich. Isopropanol and ethanol were purchased from VWR. Glycogen (Roche) was purchased from Krackeler Scientific. Random hexamers were purchased from Thermo Fisher Scientific. dNTP was purchased from BioBasic. MMLV reverse transcriptase was purchased from Clontech, and PerfeCta SYBR Green SuperMix, Low ROX, were obtained from Quanta Biosciences.

### Primers for qPCR

EHEC *tir* forward (GAAGTCGGCACCTGCGAATCA) and reverse (GCATAGGGACCGTGCAGAATC), *eae* (intimin) forward (TGTCGCACTAACAGTCGCTT) and reverse (GCAACCACGGGAAATGATGG), *espF* forward (TTCACCGGAGTAAGACGCAC) and reverse (CTGCTTCTACACTAGGGCGG), *tccp* forward (TAGCTCCATCAGCGCAACAA) and reverse (GCGCTGCCTCACATTAGGA), and *rpoA* forward (GCGCTCATCTTCTTCCGAAT) and reverse (CGCGGTCGTGGTTATGTG) primers were purchased from IDT. *C. rodentium tir* forward (CAGGCTAAACGTCAGCAGGA) and reverse (TCGGCGGATTTCGTCTATGG), *eae* (intimin) forward (TCAGCATAGCGGAAGCCAAA) and reverse (TGCTACCGCCTTGCACATAA), *espF* forward (AATGGAATTGGTCAGGCCGT) and reverse (ACTGAAAAGCTCGCACCTCC), and *rpoA* forward (GCCCTGTTGACGATCTGGAA) and reverse (GCTCAACCTCAGTACGCTGT) primers were purchased from IDT. Mouse *Muc2* forward (AATACCTGGAAGCCCAAT) and reverse (GAACACCAGTGCTCAGCGTA), *Reg3y* forward (AGTGGTAACAGTGGCCAATATGTA) and reverse (ACCACAGTGATTGCCTGAGGAAGA), *Cxcl1* forward (TGGCTGGGATTCACCTCAAG) and reverse (CCGTTACTTGGGGACACCTT), *Cxcl2* forward (CACTCTCAAGGGCGGTCAAA) and reverse (GGTTCTTCCGTTGAGGGACA), and *Rpl13a* forward (GCTGCCGAAGATGGCGGAGG) and reverse (ACCACCACCTTCCGGCCCA) were also purchased from IDT.

### ELISA

Anti-mouse TNFα (TN3-19.12, 14-7423-85), anti-mouse TNFα biotin conjugate (13-7341-81), anti-mouse IFNγ (XMG1.2, 14-7311-81), anti-mouse IFNγ biotin conjugate (R4-6A2, 13-7312-81), anti-mouse IL-17A (eBio17CK15A5, 14-7175-81), anti-mouse IL-17A biotin conjugate (eBio17B7, 13-7177-81), anti-mouse IL-1β (B122, 14-7012-81), anti-mouse IL-1β biotin conjugate (13-7112-81), and anti-mouse IL-6 (MP5-20F3, 14-7061-85) were purchased from Thermo Fisher Scientific. Anti-mouse IL-6 biotin conjugate (MP5-32C11, 554402), anti-mouse IL-12 (p40/p70) (C15.6, 551219), anti-mouse IL-12 (p40/p70) biotin conjugate (C17.8, 554476), HRP Streptavidin (554066), and TMB Substrate Reagent Set were purchased from BD Biosciences.

### Flow cytometry

Collagenase Type VII from *Clostridium histolyticum* was purchased from Sigma Aldrich, and DNase I was obtained from Roche. Percoll was obtained from Cytiva, and ACK lysis buffer was obtained from Quality Biological Inc. PE-SiglecF (Clone E50-2440, 562068) was purchased from BD Biosciences. Cell strainers (70 μm), APC-CD4 (GK1.5, 17-0041-82), APC-CD11c (N418, 17-0114-82), PerCP-Cy5.5-CD11b (M1/70, 45-0112-82), Alexa Fluor 488-TNFα (MP6-XT22, 53-7321-82), PE-IFNγ (XMG1.2, 12-7311-81), PE-IL-17A (eBio17B7, 12-7177-81), intracellular fixation buffer and permeabilization buffer were purchased from Thermo Fisher Scientific. Brefeldin A (BFA), phorbol 12-myristate 13-acetate (PMA), and ionomycin were purchased from Cayman Chemical.

### Mice

C57Bl/6, *Drd2*fl/fl (#020631, Drd2loxP), Villin-Cre (#021504), Cd11c-Cre (#008068), LysM-Cre (#004781), and Cd4-Cre (#022071) mice were acquired from Jackson Laboratories. All mice were subsequently bred and maintained at the animal facility of Cornell University and used at 8-12 weeks of age in accordance with the guidelines of the Institutional Animal Care and Use Committee and the Cornell Center for Animal Resources and Education (Protocol number 2015-0069). Mice were co-housed for 7 d prior to use and fed Envigo Teklad global irradiated 18% protein rodent diet meal 2918 as the conventional diet.

### Plasmids

DRD2-Tango (Plasmid #66269), pCDNA3.1(+)-CMV-bArrestin2-TEV (Plasmid #107245), and lentiCRISPRv2 puro (Plasmid #98290) were obtained from Addgene. pCDNA3.1-DRD2 was generated by subcloning *Drd2* into the pCDNA3.1 backbone using EcoRI and XhoI. VSVg and PAX2 packaging plasmids were obtained from Jeremy Baskin (Cornell University).

### In vivo experiments

#### Antibiotic treatment

Mice were administered an antibiotic cocktail of ampicillin (9 mg/kg), metronidazole (9 mg/kg), neomycin (9 mg/kg), and vancomycin (4.5 mg/kg) via oral gavage every 12 h. Mice were pre-treated with antibiotics for 7 d prior to receiving tryptophan diet, metabolites, or DA. Mice were continued to be administered the antibiotic cocktail while receiving tryptophan diet, metabolites, or DA. Neomycin treatment was discontinued 24 h prior to *C. rodentium* infection, and ampicillin, metronidazole, and vancomycin were continued to be administered for the remainder of the experiment.

#### Tryptophan diet

Conventional diet (Envigo Teklad global irradiated 18% protein rodent diet meal 2918) or tryptophan (Trp) diet was provided to mice *ad libitum* in feeding jars. The tryptophan diet was prepared by supplementing conventional diet (2 g Trp/kg diet) with an additional 40 g Trp/kg diet. Mice received tryptophan diet for 7 d prior to *C. rodentium* infection and then for the remainder of the experiment.

#### Metabolite treatment

Mice were administered I3A (1000 mg/kg), IPyA (2900 mg/kg), or IEt (600 mg/kg). I3A, IPyA, and IEt were dissolved in vehicle (dimethylsulfoxide, DMSO) and administered via oral gavage every 12 h. Mice were pre-treated with metabolite or vehicle for 2 d prior to *C. rodentium* infection and then for the remainder of the experiment by oral gavage.

#### Dopamine treatment

Mice were pre-treated with DA (100 mg/kg) or vehicle (PBS) via intraperitoneal (IP) injection every 24 h for 2 d prior to *C. rodentium* infection and for the remainder of the experiment.

#### DRD2 agonist treatment

Mice were pre-treated with sumanirole (SUM, 4 mg/kg) or vehicle (PBS) via IP injection every 24 h for 2 d prior to *C. rodentium* infection and for the remainder of the experiment.

#### DRD2 antagonist treatment

Mice were pre-treated with L-741,626 (1 mg/kg) or vehicle (PBS) via IP injection every 24 h for 2 d prior to Trp diet and for the remainder of the experiment.

#### SERT inhibitor treatment

Mice were pre-treated with Prozac (20 mg/kg) or vehicle (water) via oral gavage every 24 h for 2 d prior to Trp diet and for the remainder of the experiment.

#### TpH1 inhibitor treatment

Mice were pre-treated with 4-chloro-DL-phenylalanine methyl ester hydrochloride (PCPA, 150 mg/kg) or vehicle (PBS) via IP injection every 24 h for 2 d prior to Trp diet and for the remainder of the experiment.

#### C. rodentium infection

*C. rodentium* was grown in LB broth overnight with shaking at 37 °C. The next day, bacteria were subcultured until mid-log (OD600 = 0.4 – 0.6) phase and mice were infected with 10^8^ CFU by oral gavage. Mice were euthanized 10 d or 24 d post-infection.

#### Weight loss

Mice were weighed daily at the same time each day. Each mouse weight was normalized to itself and control mice on day 0.

#### CFU quantification

Feces and colon tissue were weighed and homogenized using a Corning LSE Vortex Mixer, and a Brinkmann KINEMATICA Ch-6010 KRIENS-LU Benchtop Homogenizer, respectively. Homogenates were plated onto MacConkey agar and incubated for 24 h at 37 °C. *C. rodentium* colonies were counted to determine CFU/g of feces and colon tissue.

#### Histopathology and crypt height measurement

The distal colon (2 cm segment) was flushed with PBS, excised, and fixed in 10% neutral buffered formalin, paraffin-embedded, sectioned (5 μm), and stained with Harris hematoxylin and eosin Y. Samples were blinded, imaged using an Olympus CX41RF microscope. Tissue sections were assessed for submucosal edema (0 = no change; 1 = mild; 2 = moderate; 3 = profound), goblet cell depletion (scored based on numbers of goblet cells/high-power field where 0 = > 50; 1 = 25–50; 2 = 10–25; 3 = < 10), epithelial hyperplasia (scored based on percentage above the height of the control where 0 = no change; 1 = 1–50%; 2 = 51–100%; 3 = > 100%), epithelial integrity (0 = no change; 1 = < 10 epithelial cells shedding per lesion; 2 = 11–20 epithelial cells shedding per lesion; 3 = epithelial ulceration; 4 = epithelial ulceration with severe crypt destruction) and neutrophil and mononuclear cell infiltration (0 = none; 1 = mild; 2 = moderate; 3 = severe). Data are expressed as the sum of these individual scores (0-16). Crypt height was measured using Olympus cellSens Entry software by measuring ten well-oriented crypts per mouse.

#### Cryosection immunofluorescence preparation

A portion of the distal colon was frozen in OCT, and blocks were sectioned (5 μm) on a Thermo Scientific Microm HM 525 cryostat, adhered to a glass slide, washed with PBS, and fixed with 4% PFA prior to staining.

#### Pedestal formation

Using distal colon cryosections stained with DAPI Prolong Diamond and Alexa Fluor 647-phalloidin, bacteria-associated actin pedestals were visualized. The number of pedestals per host cell were quantified for each of the three replicate images per mouse. For box plots, interquartile ranges (IQRs, boxes), median values (line within box), whiskers (lowest and highest values within 1.5 times IQR from the first and third quartiles), and outliers beyond whiskers (dots), are shown.

#### Western blot lysates

Epithelial cells from the distal colon were isolated by mechanical disruption and lysed with 1X RIPA lysis buffer (150 mM sodium chloride, 1.0% Triton X-100, 0.5% sodium deoxycholate, 0.1% SDS, 25 mM Tris, 1 mM EDTA) with 1X protease inhibitor (cOmplete tablets) and sodium β-glycerophosphate (17.5 mM), sodium orthovanadate (1 mM), sodium fluoride (20 mM), and sodium pyrophosphate decahydrate (5 mM) added immediately prior to use. Samples were quantified using the DC Protein Assay kit.

#### Griess assay

A portion of the distal colon was incubated in RPMI-1640 supplemented with 10% FBS, P/S, L-glutamine, HEPES, sodium pyruvate, and 2-mercaptoethanol at 37 °C for 24 h, after which the culture supernatant was collected. Nitric oxide (NO) production was quantified by addition of 1% (w/v) sulfanilamide in 2.5% (v/v) phosphoric acid and 0.1% (w/v) naphthylethylenediamine dihydrochloride in 2.5% (v/v) phosphoric acid. A standard curve was generated using NaNO_2_ and UV absorbance was measured at 550 nm using a BioTek PowerWave XS2 plate reader.

#### ELISA

A portion of the distal colon was incubated in RPMI-1640 supplemented with 10% FBS, P/S, L-glutamine, HEPES, sodium pyruvate, and 2-mercaptoethanol at 37 °C for 24 h, after which the culture supernatant was collected. A sandwich ELISA was performed using the following capture antibodies: anti-TNFα (1:500), anti-IFNγ (1:500), anti-IL-17A (1:1000), anti-IL-1β (1:500), anti-IL-6 (1:500), or anti-IL-12p40 (1:500). Detection was performed using the following biotin conjugated antibodies: anti-TNFα (1:500), anti-IFNγ (1:1000), anti-IL-17A (1:1000), anti-IL-1β (1:500), anti-IL-6 (1:1000), and anti-IL-12p40 (1:500). ELISAs were developed with TMB, and UV absorbance was measured at 450 nm using a BioTek PowerWave XS2 plate reader.

#### RNA isolation and qPCR

Intestinal epithelial cells were collected from the distal colon as above, and RNA was purified using RNABee according to the manufacturer’s instructions and quantified using a GE Nanovue. Using a BioRad C1000 Touch Thermal Cycler, RNA was reverse transcribed using an oligo(dT) primer and MMLV reverse transcriptase. cDNA samples were analyzed using PerfeCta SYBR Green SuperMix, Low ROX, and a BioRad CFX96 Real-Time PCR Detection System. PCR amplification conditions were as follows: 95 °C (3 min) and 40 cycles of 95 °C (15 s) and 60 °C (45 s). Relative expression of mRNA transcripts was normalized to the ribosomal protein L13a (*Rpl13a*). Data are represented as the fold induction over control samples.

#### Flow cytometry

To isolate the lamina propria from the colon, the tissue was incubated in PBS supplemented with 5% FBS, P/S, and 5 mM EDTA, shaking at 37 °C for 20 min. The colon section was then cut into small pieces and incubated in RPMI-1640 supplemented with 5% FBS, P/S, collagenase VII, and DNAse I, shaking at 37 °C for 40 min. Samples were filtered using a cell strainer, washed once, and resuspended in 8 mL of 44% Percoll and underlaid with 5 mL of 67.5% Percoll to isolate the cells. Mesenteric lymph nodes and spleens were homogenized, filtered using a cell strainer, and treated with ACK lysis buffer. For CD4^+^ T cell FACS analysis, samples were re-stimulated with PMA (50 ng/mL), ionomycin (750 ng/mL), and BFA (1:1000) for 4 h at 4 °C prior to FACS staining.

#### LC-MS

Fecal contents were collected fresh from mice and immediately flash frozen in liquid nitrogen. Frozen samples were dried on a VirTis Benchtop K Series Manifold Freeze Dryer. Dried samples were crushed and resolubilized in methanol (10x the volume of the dry weight of the samples) and rocked at room temperature for 1 h before collecting the supernatant, which was then dried down. Immediately prior to LC-MS analysis, the samples were resuspended in methanol (10x the volume of the dry weight of the samples) and filtered. LC-MS analysis was performed on an Agilent 6230 electrospray ionization–time-of-flight (ESI–TOF) MS coupled to an Agilent 1260 HPLC equipped with an Agilent Poroshell 120 ECC18 reverse phase column (3 x 50 mm, 2.7 µm) using a flow rate of 0.5 mL/min. The gradient was ramped from 90% water and 0.1% FA (Solvent A) and 10% acetonitrile and 0.1% FA (Solvent B) to 50% A and 50% B for 0.5 min. The gradient was then ramped to 35% A and 65% B for an additional 0.5 min, then to 15% A and 85% B for 4.5 min, followed by 0% A and 100% B for 0.75 min. The gradient was then held constant at 0% A and 100% B for an additional minute. For detection, the MS was equipped with a dual ESI source operating in positive or negative mode, acquiring in extended dynamic range from m/z 100–3200 at one spectrum per s; gas temperature: 325 °C; drying gas 10 L/min; nebulizer: 20 psi; fragmentor: 80 V. Quantification of metabolites was determined by integrating the extracted ion count of the exact masses of the metabolites, which were determined using commercial standards. Standard curves in which known amounts of metabolite were utilized to determine the amount of each metabolite in each sample, which was normalized to the dry weight of the fecal samples.

#### 16S rRNA gene sequencing and analysis

Following 7 d of Trp diet or 2 d of Trp metabolite treatment, genomic DNA was extracted from mouse feces using a QIAGEN DNeasy 96 PowerSoil Pro Kit. Amplicon libraries were created by PCR using universal bacterial primers targeting the V4 region of the 16S rRNA gene (515F and 806R)^44^ and Classic++ Hot Start Taq DNA Polymerase Master Mix (TONBO Biosciences). Samples were amplified in duplicate using an Applied Biosystems SimpliAmp Thermal Cycler and an Eppendorf Mastercycler Gradient Thermal Cycler. PCR amplification conditions were as follows: 94 °C (3 min), 25 cycles of 94 °C (45 s), 50 °C (1 min), 72 °C (1.5 min), and 72 °C (10 min). The duplicate amplified products were pooled, and16S rRNA gene amplicons were purified with Mag-Bind TotalPure NGS (Omega Bio-tek, Inc.). Paired-end sequencing (2×250bp) was performed on the Illumina MiSeq platform at the Cornell Institute of Biotechnology.

16S rRNA gene sequences were imported into Quantitative Insights into Microbial Ecology (QIIME2) for analysis^45^. Sequences were demultiplexed, trimmed to 250 bp, and denoised using DADA2^46^. Sequences were clustered into de novo operational taxonomic units (OTUs) with 97% similarity using the Greengenes database (version 13.8)^47^. Alpha and beta diversity analyses were computed using the ‘core-metrics-phylogenetic’ function and Shannon diversity plots and weighted UniFrac distance matrices were generated using Vega Editor (QIIME2) to generate principal coordinate analysis plots. Taxonomic classification using a pre-trained naive Bayes machine-learning classifier was performed and visualized using taxonomic bar graphs^47^.

### In vitro experiments

#### Caco-2 monolayers

Caco-2 cells were seeded at 15,000 cells per transwell insert in 12-well companion plates. Monolayers were grown in DMEM supplemented with 10% FBS and P/S at 37 °C. Media was replenished every 2-3 days. On day 18, media was replenished, and I3A, IPyA, or IEt (100 μM) or vehicle (DMSO) was added. On day 20, media and metabolites were replenished, and cells were infected with EHEC (MOI 50). After 16 h, cells were collected for Western blot or IF staining.

#### Haloperidol treatment

Caco-2 monolayers were grown as above. Haloperidol (10 μM) or vehicle (DMSO) was added initially to monolayers at on day 17 and replenished on day 18 and 20.

#### Inhibitor treatments

Caco-2 cells were seeded at 10^4^ cells/well in 12-well plates or on glass coverslips for IF. Cells were grown in DMEM supplemented with 10% FBS and P/S at 37 °C. Gαi inhibitor pertussis toxin (PTx, 100 ng/mL) was added to cells 18 h prior to metabolite treatment. β-arrestin inhibitor barbadin (100 μM), Gβγ inhibitor gallein (10 μM), PLC inhibitor U-73122 (10 μM), and PKC inhibitor sotrastaurin (5 μM) were added to cells 30 min prior to metabolite treatment. PKC-θ inhibitor (5 μM) was added to cells 24 h prior to metabolite treatment. Proteasome inhibitor MG-132 (10 μM) was added to cells 1 h prior to metabolite treatment. Following pre-treatment with inhibitor, I3A, IPyA, or IEt (100 μM) was added. After 24 h, cells were infected with EHEC (MOI 50), and 12 h post-infection, cells were collected for Western blot or IF staining.

#### PKC-θ siRNA

Control siRNA (Sense): 5’-rCrGrUrUrArArUrCrGrCrGrUrArUrArArUrArCrGrCrGrUAT-3’, (Antisense): 5’-rArUrArCrGrCrGrUrArUrUrArUrArCrGrCrGrArUrUrArArCrGrArC-3’), PKC-8 siRNA 1 (Sense): 5’-rCrArArArGrArArUrUrCrUrUrArArArCrGArGrArArGrCCC-3’, (Antisense): 5’ - rGrGrGrCrUrUrCrUrCrGrUrUrUrArArGrArArUrUrCrUrUrUrGrUrC-3’, and PKC-θ siRNA 2 (Sense): 5’-rArGrGrUrUrUrCrArArGrArCrUrUrGrArUrArCrUrGrCrAAT-3’, (Antisense): 5’-rArUrUrGrCrArGrUrArUrCrArArGrUrCrUrUrGrArArArCrCrUrUrU-3 were each incubated with Transfectagro and Lipofectamine RNAiMAX for 10 min prior to addition to Caco-2 cells. After 48 h, media was replenished, and I3A, IPyA, or IEt (100 μM) was applied to cells for 24 h, followed by infection with EHEC (MOI 50). After 12 h, cells were collected for Western blot or IF staining.

#### CRISPR/Cas9 knockout of Drd2, Drd3, and Drd4

*Drd2* forward (5′-ATGGGAGTTTCCCAGTGAAC-3′) and reverse (5′-GTTCACTGGGAAACTCCCAT-3′), *Drd3* forward (5′-CAGGCCATTGCCGAAGACGA-3′) and reverse (5′-TCGTCTTCGGCAATGGCCTG-3′), and *Drd4* forward (5′-CAACCTGTGCGCCATCAGCG-3′) and reverse (5′-CGCTGATGGCGCACAGGTTG-3′) guide RNAs were cloned into the lentiviral CRISPR plasmid lentiCRISPRv2 puro, following digestion with BsmBI. HEK 293T cells were seeded at 10^5^ cells/well in 6-well plates. At 90-95% confluency, the cells were co-transfected with the lentiCRISPR plasmid and VSVg and PAX2 packaging plasmids using Lipofectamine 2000. The cells were incubated at 37 °C for 2 d. Afterwards, the HEK 293T cell supernatant was collected and syringe-filtered using a 0.45 µm filter. For lentiviral transduction, filtered supernatant was added dropwise to Caco-2 cells seeded in a 10 cm dish, and the cells were incubated for 2 d at 37 °C. Polybrene (4 μg/mL) was added, and transduced cells were split 1:1 into media containing puromycin (2 µg/mL). After selection, successful transduction was confirmed by Western blotting.

#### Caco-2 immunofluorescence

Caco-2 monolayers or cells grown on glass coverslips were grown as above, washed with PBS, and fixed with 4% PFA prior to IF staining.

#### Pedestal formation

Using Caco-2 monolayers or cells grown on glass coverslips stained with DAPI Prolong Diamond and Alexa Fluor 647-phalloidin, bacteria-associated actin pedestals were visualized. The number of pedestals per Caco-2 cell were quantified for each of the three replicate samples per treatment. For box plots, interquartile ranges (IQRs, boxes), median values (line within box), whiskers (lowest and highest values within 1.5 times IQR from the first and third quartiles), and outliers beyond whiskers (dots), are shown.

#### Western blot lysate preparation

Caco-2 monolayers or cells were grown as above, washed with PBS, and lysed with 1X RIPA lysis buffer (150 mM sodium chloride, 1.0% Triton X-100, 0.5% sodium deoxycholate, 0.1% SDS, 25 mM Tris, 1 mM EDTA) with 1X protease inhibitor (cOmplete tablets) and sodium β-glycerophosphate (17.5 mM), sodium orthovanadate (1 mM), sodium fluoride (20 mM), and sodium pyrophosphate decahydrate (5 mM) added immediately prior to use. Samples from biological replicates were pooled prior to quantification using the DC Protein Assay kit.

#### Growth curves and viable CFU counts

EHEC or *C. rodentium* were grown in LB broth overnight with shaking at 37 °C. The next day, bacteria were cultured in fresh media with I3A, IPyA, or IEt (100 μM). Cultures were incubated with shaking at 37 °C. OD600 absorbance readings were measured after 2, 4, 6, 8, 12, and 24 h. At 24 h, cultures were plated on LB agar and incubated overnight at 37 °C. Colonies were counted to determine viable CFUs.

#### RNA isolation and qPCR

EHEC or *C. rodentium* were grown in low glucose DMEM overnight with shaking at 37 °C with I3A, IPyA, or IEt (100 μM). The next day, bacteria were subcultured until late-log phase (OD600 = 0.6 – 0.8) in low glucose DMEM with I3A, IPyA, or IEt (100 μM). Cultures were grown shaking at 250 rpm at 37 °C for aerobic conditions, standing at 37 °C for microaerophilic conditions, or standing at 37 °C under anaerobic conditions using a Coy chamber. Cells were lysed with RNABee prior to RNA purification according to the manufacturer’s instructions. RNA was quantified using a GE Nanovue. Using a BioRad C1000 Touch Thermal Cycler, RNA was reverse transcribed using random hexamers and MMLV reverse transcriptase. cDNA samples were analyzed using PerfeCta SYBR Green SuperMix, Low ROX, and a BioRad CFX96 Real-Time PCR Detection System. PCR amplification conditions were as follows: 95 °C (3 min) and 40 cycles of 95 °C (15 s) and 60 °C (45 s). Relative expression of mRNA transcripts was normalized to the RNA polymerase subunit alpha *rpoA*. Data are represented as the fold induction over control samples.

### Luciferase activity assays

Passive lysis buffer (5X PLB) was prepared with 125 mM Tris, pH 7.8, 10 mM 1,2-diaminocyclohexane tetraacetic acid (CDTA), 10 mM DTT, 5 mg/mL BSA, 5% (vol/vol) Triton X-100, and 50% (vol/vol) glycerol in ddH_2_O. An aqueous solution of 1X firefly luciferase substrate was prepared containing 75 mM HEPES, pH 8.0, 4 mM MgSO_4_, 20 mM DTT, 0.1 mM EDTA, 0.53 mM ATP, 0.27 mM coenzyme A, and 0.47 mM D-luciferin (firefly) in ddH_2_O. An aqueous solution of 1X Renilla luciferase buffer was prepared containing 7.5 mM sodium acetate, pH 5.0, 400 mM sodium sulfate, 10 mM CDTA, 15 mM sodium pyrophosphate, and 0.025 mM 2-(4-aminophenyl)-6-methylbenzothiazole. A 100X Renilla luciferase substrate was prepared by diluting coelenterazine to 0.55 mM in anhydrous methanol and added to 1X Renilla luciferase buffer immediately prior to the assay.

#### DRD2-Tango assay

HEK 293T cells were transfected overnight with DRD2-Tango and pCDNA3.1(+)-CMV-bArrestin2-TEV. The next day, media was replenished, and cells were treated for 24 h with I3A, IPyA, IET, or DA (100 μM) or vehicle (DMSO). After 24 h, cells were washed with PBS, lysed with 1X PLB, and 20 µl of lysate was added to a 96-well white flat-bottom plate. Afterwards, 50 µl of 1X firefly luciferase substrate was added to each well, and luminescence was measured for 10 min using a Turner BioSystems Veritas Microplate Luminometer. Immediately after, 50 µl of the 1X Renilla substrate was added to each well, and luminescence was measured for 10 min. Luciferase activity was determined by calculating the ratio of the firefly luciferase signal to the Renilla luciferase signal.

#### GloSensor assay

HEK 293T cells stably expressing a GloSensor cAMP reporter were transfected overnight with pCDNA3.1-DRD2. The next day, media was replenished, and the cells were incubated with D-luciferin (0.47 mM) for 30 min at room temperature. Cells were then treated for 15 min with I3A, IPyA, IEt, or DA (100 μM, unless otherwise indicated for EC50 and Kd determination). After 15 min, cells were washed with PBS, lysed with 1X PLB, and 20 µl of lysate was added to a 96-well white flat-bottom plate. Afterwards, luminescence was measured as above using a Turner BioSystems Veritas Microplate Luminometer. Kd values were calculated using GraphPad Prism.

### Western blot

Lysates were sonicated using a Heat Systems Ultrasonic Processor XL sonicator. Protein concentrations were determined using the DC Protein Assay kit and BioTek PowerWave XS2 plate reader. Lysates were resolved on SDS-polyacrylamide gels and transferred to nitrocellulose. Membranes were blocked with 5% BSA in 25 mM Tris, 150 mM sodium chloride, and 0.1% Tween-20 solution (TBS-T) rocking at room temperature for 1 h, then probed with the appropriate antibodies (anti-DRD2, DRD3, DRD4, PKC-θ, pWIP, WIP, pN-WASP, and N-WASP primary antibodies, 1:1000; anti-β-actin antibody (1:20,000); anti-α-tubulin antibody (1:10,000); and anti-GAPDH (1:2,500)) in 5% milk or BSA in TBS-T with 0.05% sodium azide with rocking at 4 °C overnight. Overnight incubation in primary antibody was followed by washing (3 x TBS-T) and incubation with the appropriate species-specific HRP antibody (1:10,000) in 5% milk in TBS-T with rocking at room temperature for 1 h. Western blots were developed using SuperSignal West Pico Chemiluminescent Substrate on a BioRad ChemiDoc MP. Densitometry was performed using FIJI and normalized to the housekeeping protein and control lysate proteins.

### Immunofluorescence

Fixed samples were permeabilized with 0.5% Triton X-100 in PBS for 15 min, blocked with 5% BSA in PBS for 1 h, and then incubated with the anti-DRD2 antibody (1:200 dilution) in 5% BSA in PBS at room temperature for 2 h. Samples were incubated with donkey anti-mouse Alexa Fluor 594 antibody (1:500) in 5% BSA in PBS in the dark at room temperature for 1 h, then mounted with DAPI Prolong Diamond overnight. Alternatively, the samples were incubated with Alexa Fluor 647-phalloidin (1:500) in 5% BSA in PBS in the dark at room temperature for 1 h, then mounted with DAPI Prolong Diamond overnight. Samples were imaged with a Zeiss LSM 800 confocal laser scanning microscope equipped with 20X 0.8 NA and 40X 1.4 NA Plan Apochromat objectives, 405, 488, 561, and 640 nm solid-state lasers, and two GaAsP PMT detectors or a Zeiss LSM 880 confocal laser scanning microscope equipped with a 40X 1.4 NA Plan Apochromat objective, 405, 458, 488, 514, 561, and 633 nm solid-state lasers, two PMT channels, and a 32 channel GaAsP detector array. Relative brightness of stained cells was quantified using FIJI and normalized to control cells.

### Similarity ensemble approach (SEA)

SEA was utilized to predict protein targets for I3A, IPyA, and IEt. SEA is provided by the Shoichet Laboratory in the Department of Pharmaceutical Chemistry at the University of California, San Francisco at sea.bkslab.org. Non-human protein targets were excluded.

### Statistical analysis

Experiments were completed at least three independent times. Error bars signify standard deviation from the mean. Unless otherwise indicated, statistical significance between two groups was evaluated with a two-tailed Student’s t-test or determined using one-way ANOVA followed by post-hoc Tukey’s test for multiple comparison analyses.

## Acknowledgements

We are grateful to the Arnold and Mabel Beckman Foundation (Beckman Young Investigator Award to P.V.C.) and a President’s Council for Cornell Women Affinito-Stewart Grant (P.V.C.) for support. This work was supported in part by a grant from the National Institutes of Health (NIH R35GM133501). J.F. was supported by a Cornell Institute of Host-Microbe Interactions & Disease (CIHMID) Postdoctoral Fellowship. Imaging data was acquired through the Cornell Institute of Biotechnology’s BRC Imaging Facility (RRID:SCR_021741), with NYSTEM (C029155) and NIH (S10OD018516) funding for the shared Zeiss LSM880 confocal/multiphoton microscope. We thank the Weill Institute for Cell and Molecular Biology for additional resources and reagents.

## Author contributions

S.A.S. and P.V.C. conceptualized the project; S.A.S., J.F., P.V.C. designed and carried out the studies; S.A.S., J.F., P.V.C. wrote the manuscript.

## Data availability

All data are available in the main text or the Extended Data.

## Materials & Correspondence

Correspondence and requests for materials should be addressed to P.V.C.

