## Supplementary Information for "Dopamine receptor D2 confers colonization resistance via gut microbial metabolites"

**This PDF file includes:**

Extended Data Figs. 1 to 21

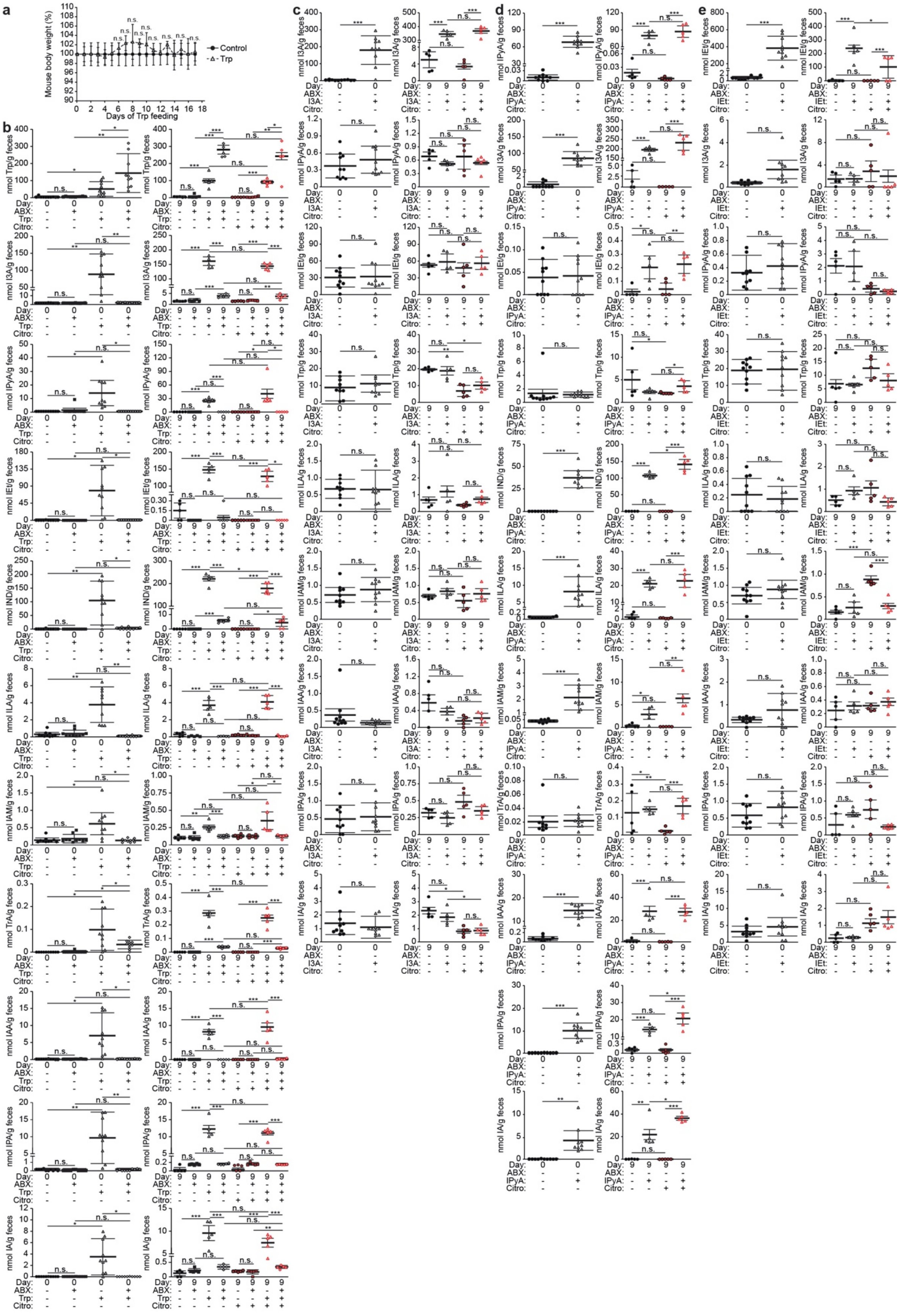
a
Mouse body weight (%)
Days of Trp feeding
Control
Trp
b
c
d
e

**Extended Data Fig. 1. LC–MS quantification of tryptophan (Trp) metabolites during Trp diet or metabolite treatment in mouse model of EHEC infection with *Citrobacter rodentium*, strain DBS100.** C57Bl/6 mice were pre-treated with antibiotics (-/+ ABX, 9 mg/kg each of metronidazole, ampicillin, and neomycin, 4.5 mg/kg of vancomycin) for 7 d, followed by conventional (2 g Trp/kg diet, ad libitum) or **a, b,** Trp (42 g Trp/kg diet, ad libitum) diet for 7 d or **c–e**, Trp metabolites, (**c**) I3A (1000 mg/kg), (**d**) IPyA (2900 mg/kg), or (**e**) IEt (600mg/kg), by oral gavage daily for 2 d. The mice were then administered *C. rodentium* (CR, oral gavage, $10^8$ colony-forming units, CFU) with continued ABX (except neomycin) and (**a, b**) Trp feeding or (**c–e**) metabolite treatment. (**a**) Mice were weighed daily during Trp diet. (**b-e**) Feces were collected before infection and euthanasia to determine Trp metabolite levels. Metabolites from the fecal colonic contents were measured by mass spectrometry, using commercial standards for quantification. Abbreviations: Indole-3-aldehyde (I3A), indole-3-pyruvate (IPyA), tryptophol (IEt), indole (IND), indole-3-acetamide (IAM), DL-indole-3-lactate (ILA), indole-3-acetic acid (IAA), indole-3-propionate (IPA), tryptamine hydrochloride (TrA), indole-3-acrylate (IA). Data are representative of at least 3 independent experiments, n = 5-10 mice per group. Two-tailed Student's t-test or one-way ANOVA followed by post-hoc Tukey's test: *p<0.05, **p<0.01, ***p<0.001, n.s. = not significant.

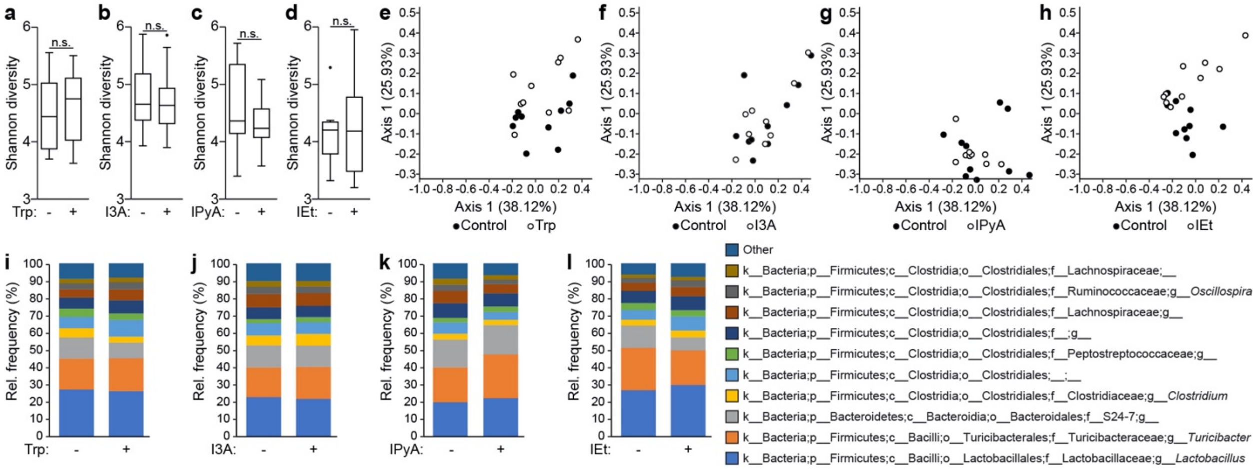


**Extended Data Fig. 2. Trp diet and metabolites do not affect gut microbial composition.** C57Bl/6 mice were treated with **a, e, i,** Trp (42 g Trp/kg diet, ad libitum) or conventional (2 g Trp/kg diet, ad libitum) diet for 7 d or (**b–d, f–h, j–l**) Trp metabolites (I3A, 1000 mg/kg; IPyA, 2900 mg/kg; or IEt, 600 mg/kg) for 2 d. Fecal samples were collected and DNA extracted, followed by 16S rRNA gene analysis. (**a–d**) Alpha diversity was measured by the Shannon diversity index. (**e–h**) Beta diversity was measured by weighted UniFrac and plotted using principal component analysis. Permutational multivariate analysis of variance (PERMANOVA) was used to assess differences in beta diversity. (**i–l**) Microbial composition is represented by taxonomic bar graphs. Data are representative of at least 3 independent experiments, n = 10 mice per group. Two-tailed Student's t-test: n.s. = not significant.

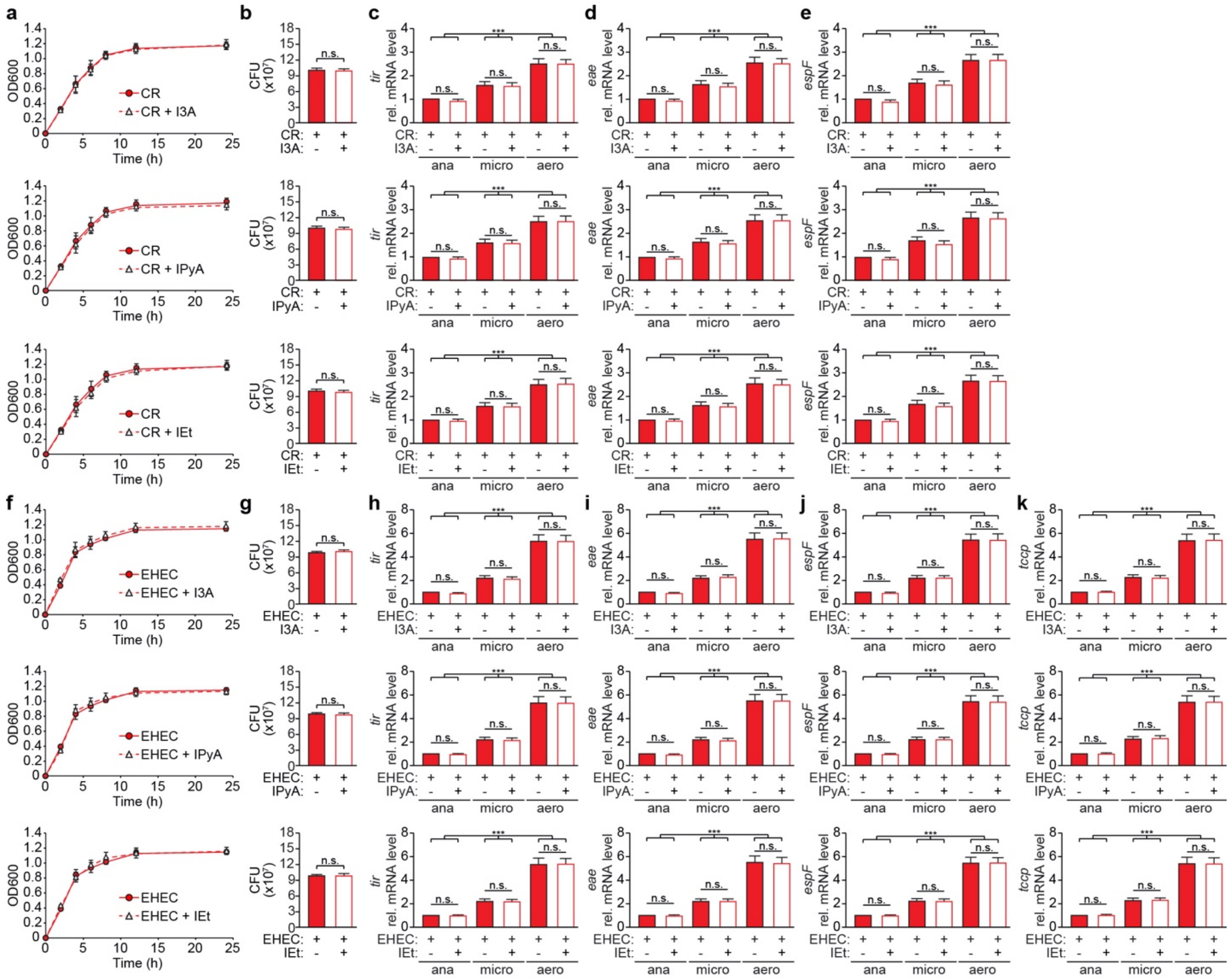


**Extended Data Fig. 3. Tryptophan metabolites do not affect *C. rodentium* and EHEC growth and virulence in vitro.** *C. rodentium* (CR) (**a–e**) and EHEC O157:H7 (**f–k**) were cultured in the presence of I3A, IPyA, or IEt (100 µM). **a, f**, Growth was monitored by measuring OD600 absorbance readings over 24 h. **b, g**, Cultures were plated after 24 h, and CFUs were counted. (**c–e, h–k**) Bacteria were cultured in low glucose DMEM under anaerobic (ana), microaerophilic (micro), and aerobic (aero) conditions to activate locus of enterocyte effacement-pathogenicity island expression with I3A, IPyA, or IEt (100 µM). RNA was isolated after cultures reached late-log phase (OD600 = 0.6–0.8), and cDNA was synthesized and analyzed by qPCR for the indicated genes. Relative expression of mRNA transcripts was normalized to the RNA polymerase subunit alpha *rpoA*. Data are represented as the fold induction over control samples. Two-tailed Student's t-test or one-way ANOVA followed by post-hoc Tukey's test: n=3, ***$p<0.001$, all other points not indicated are n.s. = not significant.

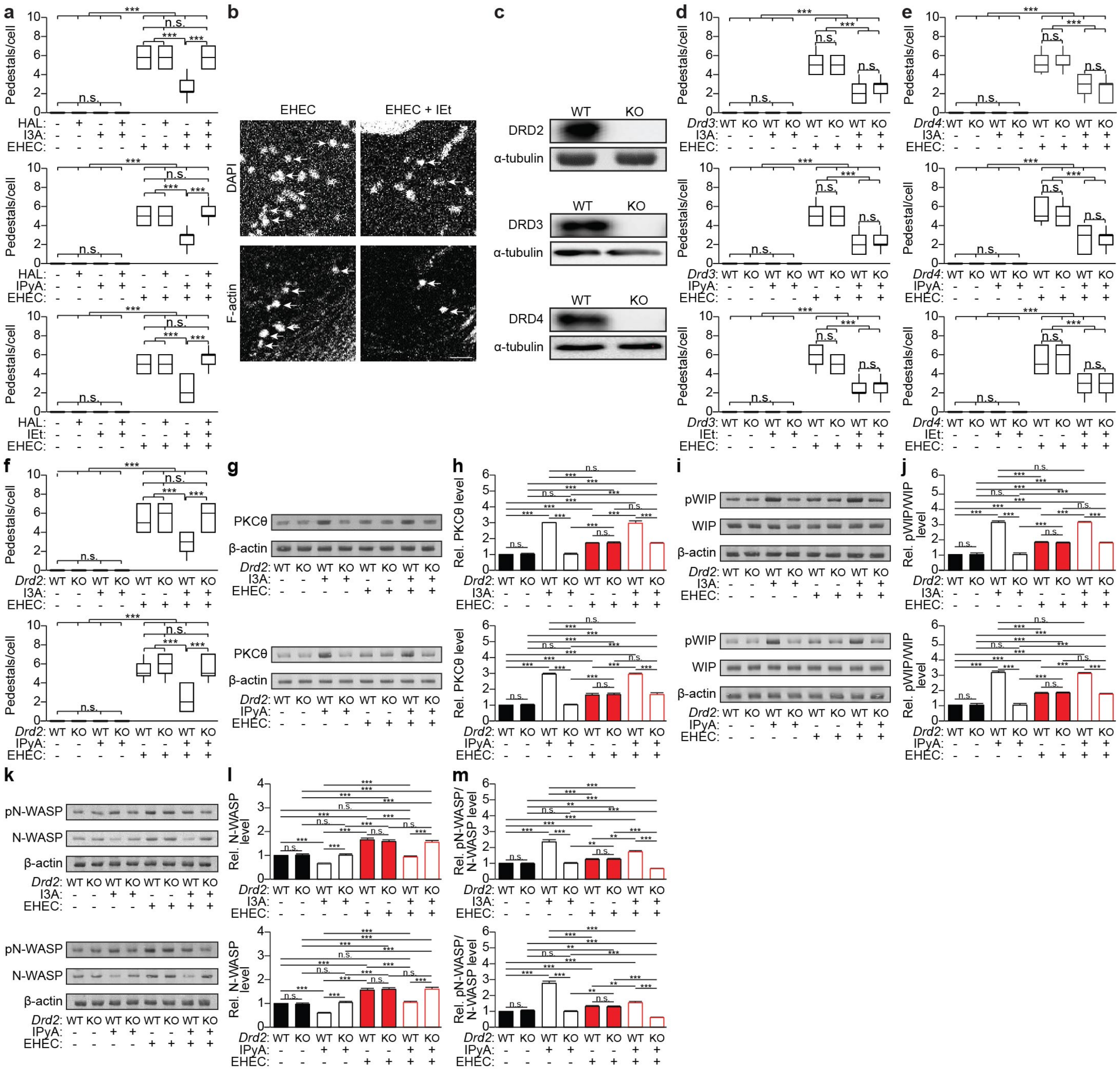


**Extended Data Fig. 4. Effects of I3A, IPyA, and IEt depend on dopamine receptor D2 (DRD2). a–b**, Polarized Caco-2 monolayers were pre-treated with haloperidol (HAL, 10 µM) for 24 h, followed by metabolites (I3A, IPyA, or IEt, 100 µM) for 2 d, and then infection with EHEC O157:H7 for 16 h. **b**, Representative images of pedestals (denoted by arrows) from Caco-2 cells stained with DAPI and Alexa Fluor 647-phalloidin and imaged by confocal microscopy. Shown are maximum intensity z-projections. Scale bar: 5 µm. **c**, Western blot analysis of Caco-2 monolayers to verify CRISPR/Cas9-mediated knockout (KO) of *Drd2*, *Drd3*, and *Drd4*. α-tubulin is shown as a loading control. **d–f**, Caco-2 monolayers (WT vs. KO) were pre-treated with metabolites (I3A, IPyA, or IEt, 100 µM) for 2 d and then infected with EHEC O157:H7 for 16 h. (**a, d–f**) Pedestal formation = # of pedestals per Caco-2 cell (HAL: I3A, n = 135–146; IPyA, n = 114–181; IEt, n = 157–192; *Drd3* KO: I3A, n = 153–188; IPyA, n = 151–185; IEt, n = 125–169; *Drd4* KO: I3A, n = 150–187; IPyA, n = 139–157; IEt, n = 101–138; *Drd2* KO: I3A, n = 109–182;

IPyA, n = 116–177; IEt, see Fig. 4c). **g-m**, Caco-2 cells (WT vs KO) were pre-treated with metabolites (I3A, IPyA, or IEt, 100 µM) for 24 h and then infected with EHEC O157:H7 for 12 h. Cell lysates were analyzed by Western blotting with the indicated antibodies. (**h, j, l, m**) Densitometry was performed using FIJI. One-way ANOVA followed by post-hoc Tukey's test: **p< 0.01, ***p<0.001, n.s. = not significant.

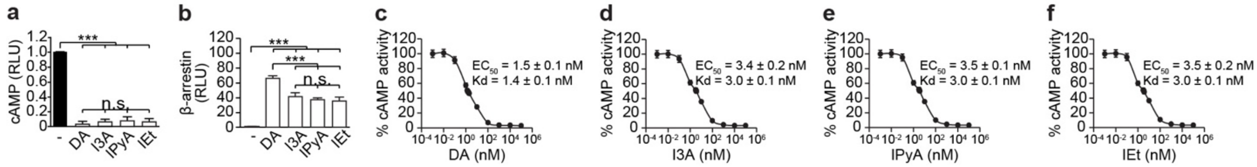


**Extended Data Fig. 5. I3A, IPyA, and IEt are ligands of dopamine receptor D2 (DRD2), which signals via Gαi. a, c–f**, HEK 293T cells overexpressing either DRD2 and a split luciferase-based cAMP sensor (GloSensor) or **b**, DRD2-Tango and a β-arrestin-TEV fusion were incubated with dopamine (DA), I3A, IPyA, or IEt (1 mM each in **a–b**; concentrations indicated in **c–f**) for 15 min (**a, c–f**) or 24 h (**b**), after which luminescence was measured to quantify ligand-induced (**a**) decrease in cAMP or (**b**) increase in β-arrestin recruitment. RLU = relative luminescence units. Kd values were calculated using GraphPad Prism. One-way ANOVA followed by post-hoc Tukey's test: ***p<0.001, n.s. = not significant.

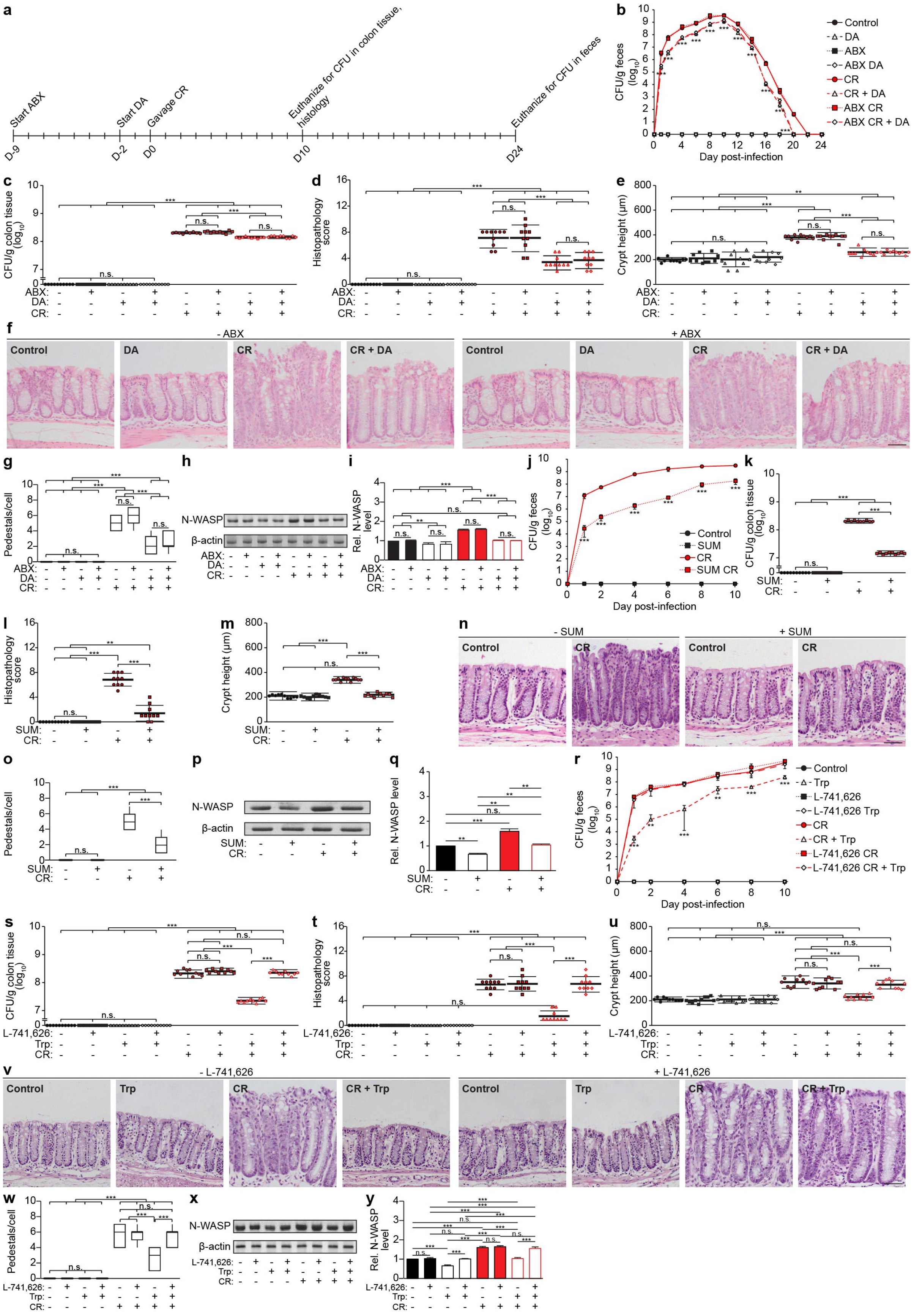
a
Start ABX
Start DA
Gavage CR
Euthanize for CFU in colon tissue, histology
Euthanize for CFU in feces
D-9
D-2
D0
D10
D24
b
CFU/g feces (log10)
Day post-infection
Control
DA
ABX
ABX DA
CR
CR + DA
ABX CR
ABX CR + DA
c
CFU/g colon tissue (log10)
ABX:
DA:
CR:
n.s.
d
Histopathology score
e
Crypt height (μm)
f
- ABX
+ ABX
Control
DA
CR
CR + DA
g
Pedestals/cell
h
N-WASP
β-actin
i
Rel. N-WASP level
j
Control
SUM
CR
SUM CR
Day post-infection
k
SUM:
l
m
n
- SUM
+ SUM
o
p
q
r
Control
Trp
L-741,626
L-741,626 Trp
CR
CR + Trp
L-741,626 CR
L-741,626 CR + Trp
s
L-741,626:
Trp:
t
u
v
- L-741,626
+ L-741,626
w
x
y

**Extended Data Fig. 6. Dopamine and a synthetic DRD2 agonist protect against a mouse model of EHEC infection using *C. rodentium*, strain DBS100, whereas a DRD2 antagonist blocks the effects. a–i**, C57Bl/6 mice were pre-treated with antibiotics (-/+ ABX, 9 mg/kg each of metronidazole, ampicillin, and neomycin, 4.5 mg/kg of vancomycin) for 7 d, followed by dopamine (DA, 100 mg/kg, intraperitoneal (IP) injection) daily for 2 d. **j–q**, C57Bl/6 mice were pre-treated with DRD2 agonist sumanirole (SUM, 4 mg/kg, IP injection) daily for 2 d. **r–y**, C57Bl/6 mice were pre-treated with DRD2 antagonist L-741,626 (1 mg/kg, IP injection) daily for 2 d, followed by conventional (2 g Trp/kg diet, ad libitum) or Trp (42 g Trp/kg diet, ad libitum) diet for 7 d. **a–y**, The mice were then administered *C. rodentium* (CR, oral gavage, $10^8$ colony-forming units, CFU) with (**a–i)** continued ABX (except neomycin) and DA treatment, (**j–q**) SUM treatment, or (**r–y**) L-741,626 and Trp feeding. **a**, Timeline for DA study. **b–c, j–k, r–s**, Bacterial load in (**b, j, r**) feces and (**c, k, s**) colon tissue was measured (**b, j, r**) every 1–2 d for (**b**) 24 d or (**j, r**) 10 d post-infection and (**c, k, s**) at the peak of infection, 10 d post-infection. (**d–f, l–n, t–v**) Colon sections were stained with H&E and (**d, l, t** blindly scored for submucosal edema (0-3), goblet cell depletion (0-3), epithelial hyperplasia (0-3), epithelial integrity (0-4), and neutrophil and mononuclear cell infiltration (0-3). Data are expressed as the sum of these individual scores (0-16). See Methods for full description of scoring rubric. (**e, m, u**) Crypt heights were measured. (**f, n, v**) Representative images. Scale bar: 50 µm. (**g, o, w**) Intestinal cryosections were stained with DAPI and Alexa Fluor 647-phalloidin. Pedestal formation = # of pedestals per host cell (DA, n = 133–140; SUM, n = 104–139; L-741,626, n = 153–189). **h–i, p–q, x–y**, Intestinal epithelial cells were isolated, and cell lysates were analyzed by Western blotting with the indicated antibodies. (**i, q, y**) Densitometry was performed using FIJI. Data are representative of at least 3 independent experiments, n=10 mice per group, bars = mean, error bars = standard deviation. One-way ANOVA followed by post-hoc Tukey's test: $^{**}p<0.01$, $^{***}p<0.001$, n.s. = not significant.

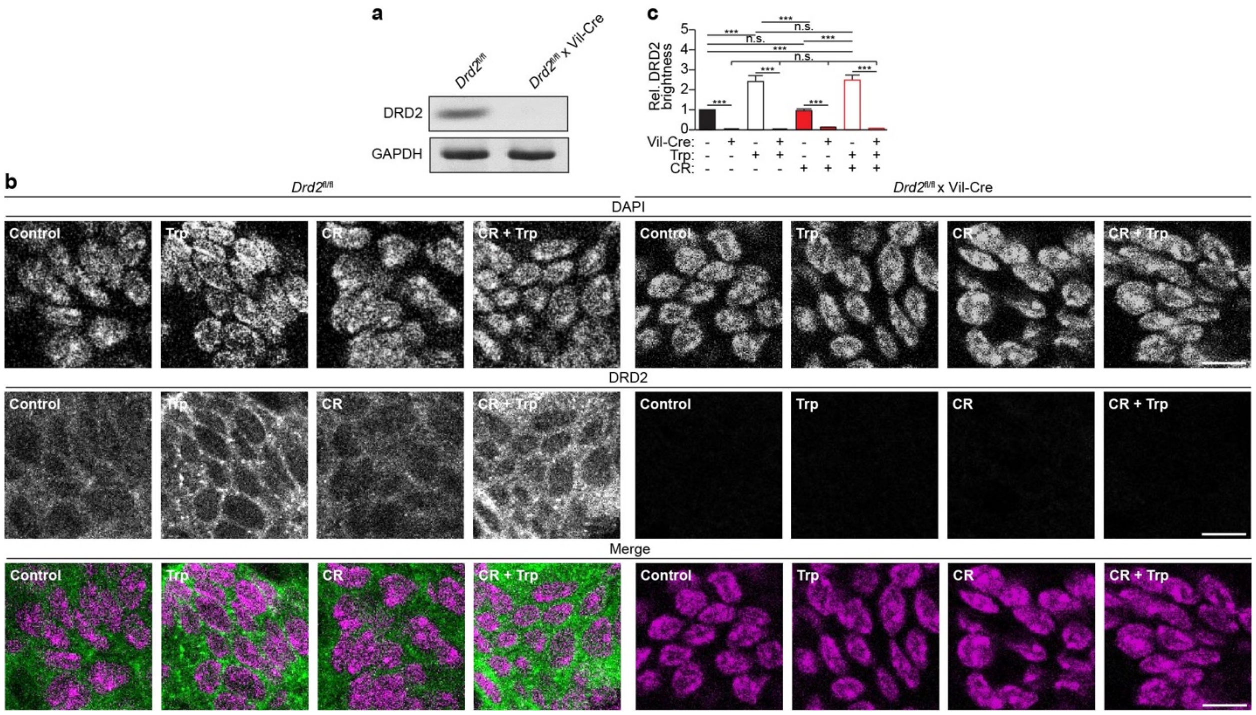


**Extended Data Fig. 7.** ***Drd2*** **is knocked out in intestinal epithelial cells in** ***Drd2*****fl/fl x Villin-Cre mice.** (**a**) Intestinal epithelial cells were isolated from *Drd2*fl/fl vs. *Drd2*fl/fl x Villin (Vil)-Cre, and cell lysates were analyzed by Western blotting with the indicated antibodies. (**b–c**) *Drd2*fl/fl x Villin (Vil)-Cre or *Drd2*fl/fl mice were fed a conventional (2 g Trp/kg diet, ad libitum) or Trp (42 g Trp/kg diet, ad libitum) diet for 7 d and then infected with *C. rodentium* (CR, oral gavage, $10^8$ CFU) with continued Trp feeding. Ten days post-infection, intestinal cryosections were stained with DAPI and an anti-DRD2 antibody, followed by an anti-mouse Alexa Fluor 594 antibody. (**b**) Shown are representative z-slices. Scale bar: 20 µm. (**c**) Image brightness was quantified using FIJI. Data are representative of at least 3 independent experiments, n=10 mice per group, bars = mean, error bars = standard deviation. One-way ANOVA followed by post-hoc Tukey's test: ***p<0.001, n.s. = not significant.

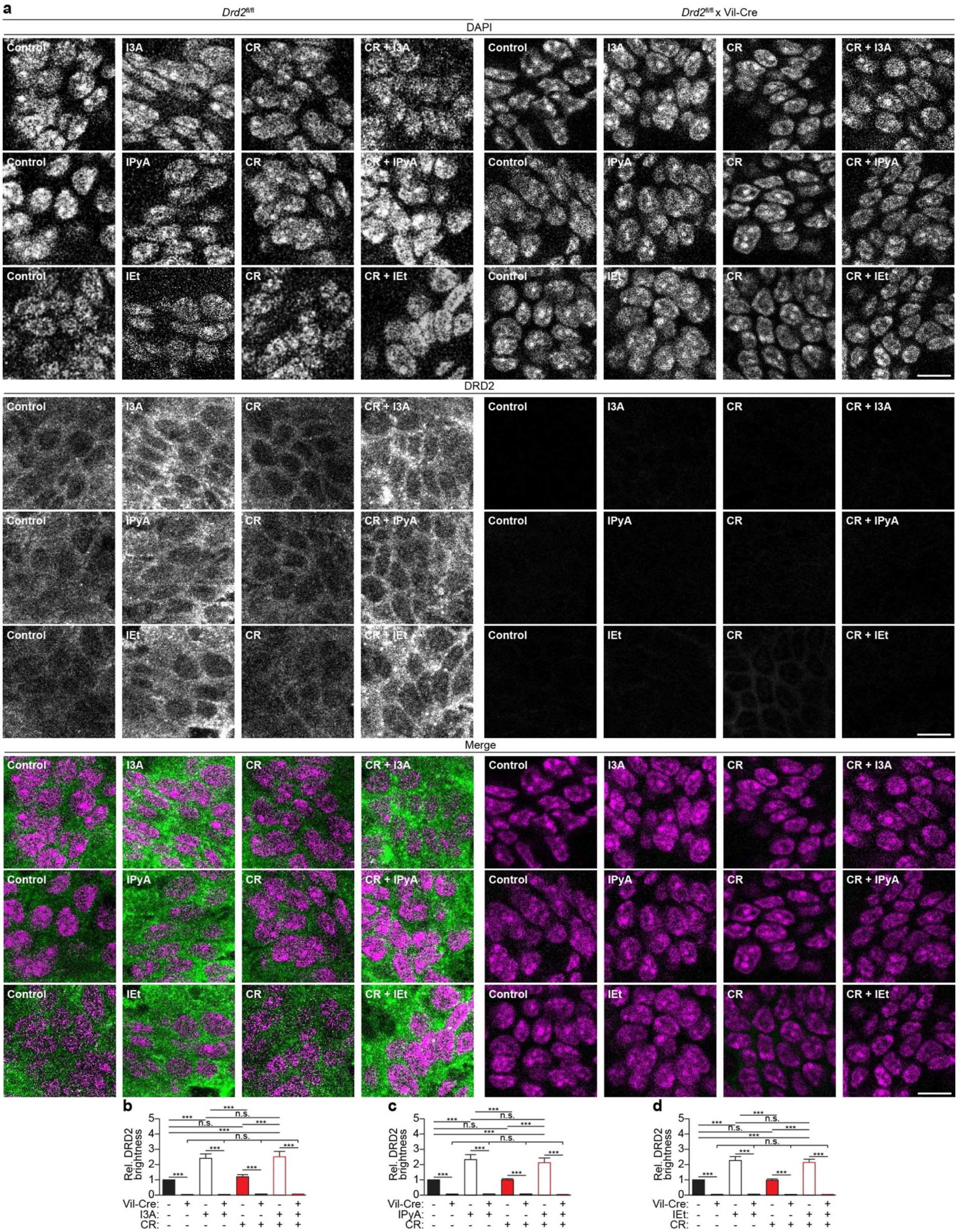
a
Drd2fl/fl
Drd2fl/fl x Vil-Cre
DAPI
DRD2
Merge
Control
I3A
CR
CR + I3A
IPyA
CR + IPyA
IEt
CR + IEt
b
c
d
Rel. DRD2 brightness
Vil-Cre:
I3A:
IPyA:
IEt:
CR:
***
n.s.

**Extended Data Fig. 8. *Drd2* is knocked out in intestinal epithelial cells in *Drd2*fl/fl x Villin-Cre mice.** *Drd2*fl/fl x Villin (Vil)-Cre or *Drd2*fl/fl mice were treated with Trp metabolites, I3A (1000 mg/kg), IPyA (2900 mg/kg), or IEt (600 mg/kg), by oral gavage daily for 2 d, and then infected with *C. rodentium* (CR, oral gavage, $10^8$ CFU) with continued metabolite treatment. Ten days post-infection, intestinal cryosections were stained with DAPI and an anti-DRD2 antibody, followed by an anti-mouse Alexa Fluor 594 antibody. (**a**) Shown are representative z-slices. Scale bar: 20 μm. (**b**–**d**) Image brightness was quantified using FIJI. Data are representative of at least 3 independent experiments, n=10 mice per group, bars = mean, error bars = standard deviation. One-way ANOVA followed by post-hoc Tukey's test: ***p<0.001, n.s. = not significant.

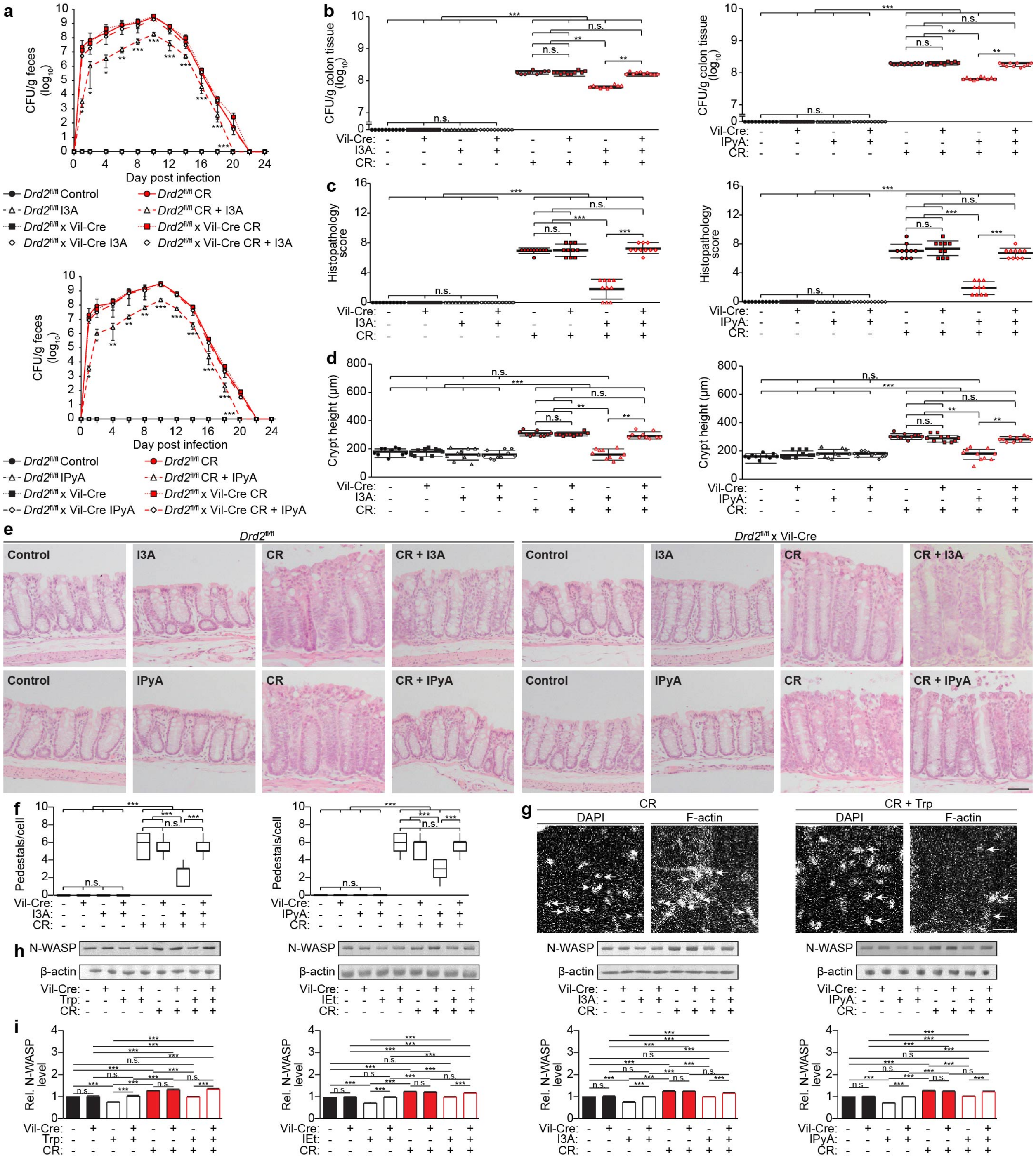

a
CFU/g feces (log10)
Day post infection
Drd2fl/fl Control
Drd2fl/fl CR
Drd2fl/fl I3A
Drd2fl/fl CR + I3A
Drd2fl/fl x Vil-Cre
Drd2fl/fl x Vil-Cre CR
Drd2fl/fl x Vil-Cre I3A
Drd2fl/fl x Vil-Cre CR + I3A
Drd2fl/fl IPyA
Drd2fl/fl CR + IPyA
Drd2fl/fl x Vil-Cre IPyA
Drd2fl/fl x Vil-Cre CR + IPyA
b
CFU/g colon tissue (log10)
c
Histopathology score
d
Crypt height (μm)
Vil-Cre:
I3A:
IPyA:
CR:
e
Control
I3A
CR
CR + I3A
IPyA
CR + IPyA
Drd2fl/fl x Vil-Cre
f
Pedestals/cell
g
CR
CR + Trp
DAPI
F-actin
h
N-WASP
β-actin
Trp:
IEt:
i
Rel. N-WASP level

**Extended Data Fig. 9. Effects of the tryptophan (Trp) diet and metabolites I3A, IPyA, and IEt in protecting against *C. rodentium* infection depend on dopamine receptor D2 (DRD2) in intestinal epithelial cells (IECs).** *Drd2*fl/fl x Villin (Vil)-Cre and *Drd2*fl/fl mice were fed a conventional (2 g Trp/kg diet, ad libitum) or Trp (42 g Trp/kg diet, ad libitum) diet for 7 d or Trp metabolites, I3A (1000 mg/kg), IPyA (2900 mg/kg), or IEt (600 mg/kg), by oral gavage daily for 2 d, and then infected with *C. rodentium* (CR, oral gavage, $10^8$ CFU) with continued Trp feeding or metabolite treatment. **a–b**, Bacterial load in (**a**) feces and (**b**) colon tissue was measured (**a**) every 1–2 d for 24 d post-infection and (**b**) at the peak of infection, 10 d post-infection. **c–e**, Colon sections were stained with H&E and (**c**) blindly scored for submucosal edema (0-3), goblet cell depletion (0-3), epithelial hyperplasia (0-3), epithelial integrity (0-4), and neutrophil and mononuclear cell infiltration (0-3). Data are expressed as the sum of these individual scores (0-16). See Methods for full description of scoring rubric. (**d**) Crypt heights were measured. (**e**) Representative images. Scale bar: 50 µm. **f, g**, Pedestal formation = # of pedestals per host cell (I3A, n = 163–179; IPyA, n = 101 – 183). (**g**) Representative images of pedestals from Fig. 4a (denoted by arrows) stained with DAPI and Alexa Fluor 647-phalloidin and imaged by confocal microscopy. Shown are maximum intensity z-projections. Scale bar: 5 µm. **h, i**, Intestinal epithelial cells were isolated, and cell lysates were analyzed by Western blotting with the indicated antibodies. (**i**) Densitometry was performed using FIJI. Data are representative of at least 3 independent experiments, n=10 mice per group, bars = mean, error bars = standard deviation. One-way ANOVA followed by post-hoc Tukey's test: *p<0.05, **p<0.01, ***p<0.001, n.s. = not significant.

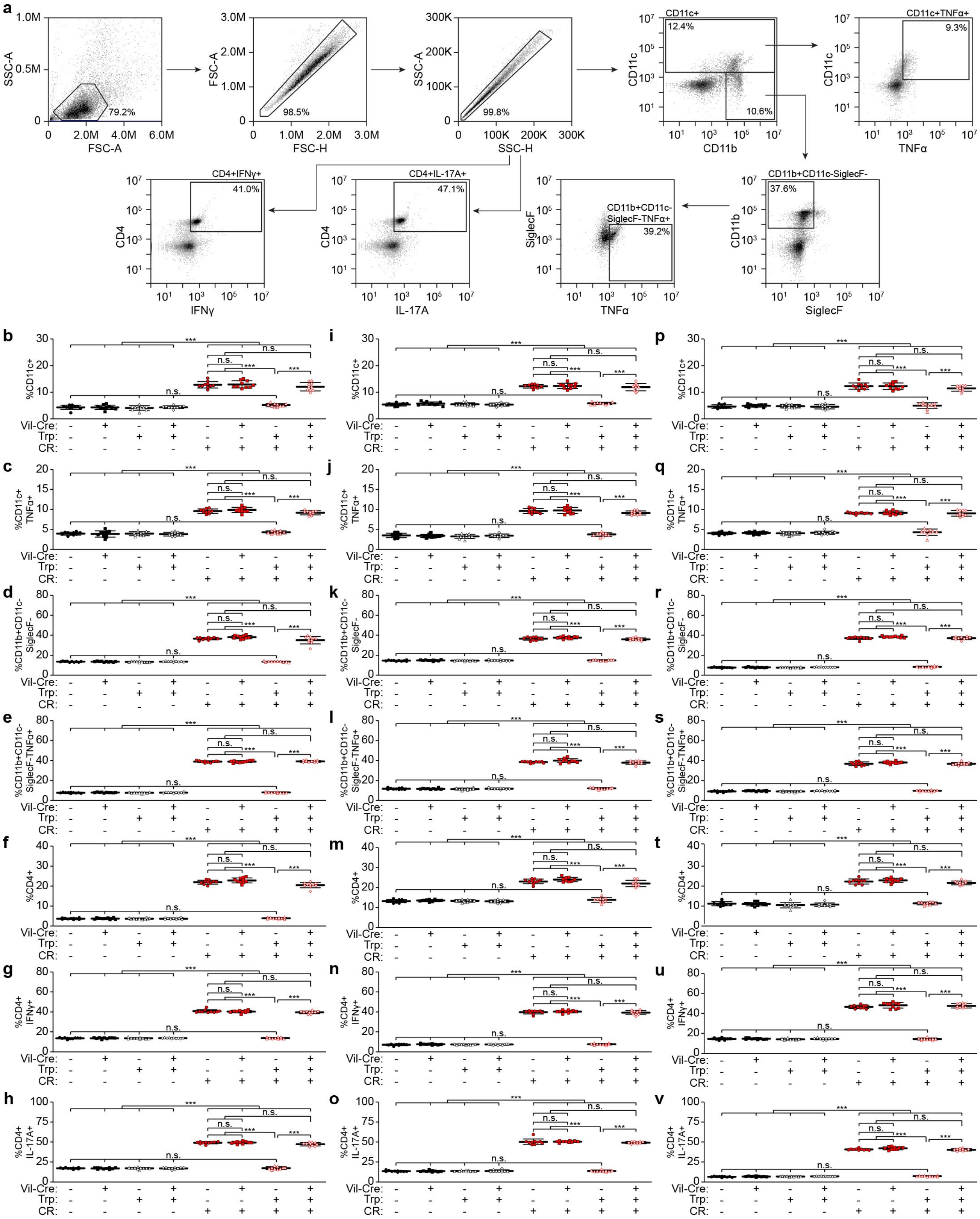
a
SSC-A
FSC-A
79.2%
FSC-H
98.5%
SSC-H
99.8%
CD11c+
12.4%
10.6%
CD11c
CD11b
CD11c+TNFα+
9.3%
TNFα
CD11b+CD11c-SiglecF-
37.6%
SiglecF
CD11b+CD11c-SiglecF-TNFα+
39.2%
CD4+IL-17A+
47.1%
IL-17A
CD4+IFNγ+
41.0%
IFNγ
CD4
b
c
d
e
f
g
h
i
j
k
l
m
n
o
p
q
r
s
t
u
v
%CD11c+
%CD11c+ TNFα+
%CD11b+CD11c- SiglecF-
%CD11b+CD11c- SiglecF-TNFα+
%CD4+
%CD4+ IFNγ+
%CD4+ IL-17A+
Vil-Cre:
Trp:
CR:
n.s.
***

**Extended Data Fig. 10. Trp diet decreases immune cell infiltrates and inflammation via DRD2 expressed in IECs.** *Drd2*fl/fl x Villin (Vil)-Cre and *Drd2*fl/fl mice were fed a conventional (2 g Trp/kg diet, ad libitum) or Trp (42 g Trp/kg diet, ad libitum) diet for 7 d, and then infected with *C. rodentium* (CR, oral gavage, $10^8$ CFU) with continued Trp feeding. **a**, Representative FACS gating strategy. **b–v**, Indicated cell populations were analyzed from the (**b–h**) lamina propria (LP), (**i–o**) mesenteric lymph nodes **(**MLN), and (**p–v**) spleen (SP) by FACS. Data are representative of at least 3 independent experiments, n=10 mice per group, bars = mean, error bars = standard deviation. One-way ANOVA followed by post-hoc Tukey's test: ***$p<0.001$, n.s. = not significant.

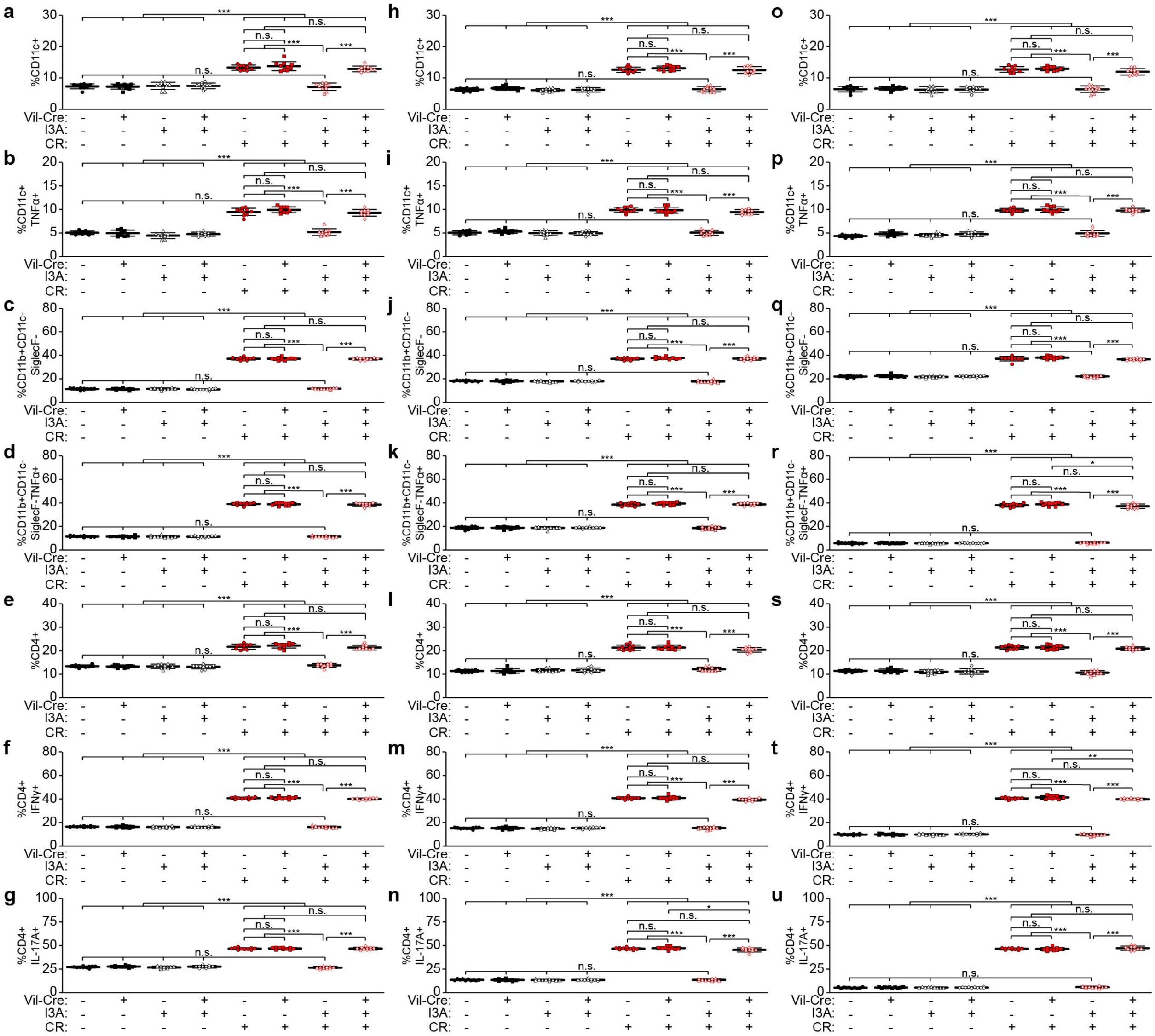


**Extended Data Fig. 11. I3A decreases immune cell infiltrates and inflammation via DRD2 expressed in IECs.** *Drd2*fl/fl x Villin (Vil)-Cre and *Drd2*fl/fl mice were administered I3A (1000 mg/kg) daily by oral gavage for 2 d, and then infected with *C. rodentium* (CR, oral gavage, $10^8$ CFU) with continued metabolite treatment. Representative gating strategy, see Extended Data Fig. 10a. **a**–**u**, Indicated cell populations were analyzed from the (**a**–**g**) LP, (**h**–**n**) MLN, and (**o**–**u**) SP by FACS. Data are representative of at least 3 independent experiments, n=10 mice per group, bars = mean, error bars = standard deviation. One-way ANOVA followed by post-hoc Tukey's test: *p<0.05, **p<0.01, ***p<0.001, n.s. = not significant.

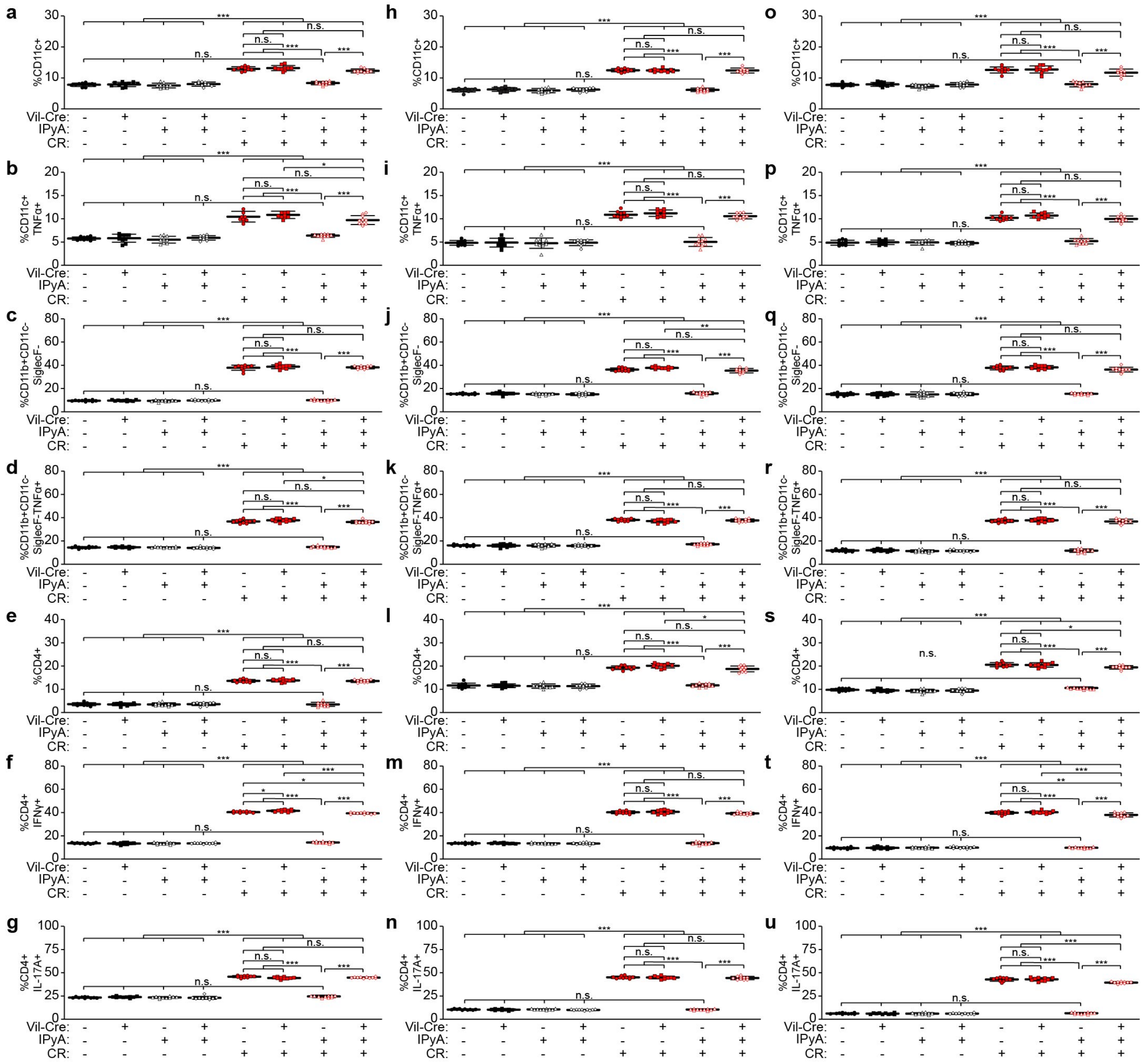

**Extended Data Fig. 12. IPyA decreases immune cell infiltrates and inflammation via DRD2 expressed in IECs.** *Drd2*fl/fl x Villin (Vil)-Cre and *Drd2*fl/fl mice were administered IPyA (2900 mg/kg) daily by oral gavage for 2 d, and then infected with *C. rodentium* (CR, oral gavage, $10^8$ CFU) with continued metabolite treatment. Representative gating strategy, see Extended Data Fig. 10a. **a**–**u**, Indicated cell populations were analyzed from the (**a**–**g**) LP, (**h**–**n**) MLN, and (**o**–**u**) SP by FACS. Data are representative of at least 3 independent experiments, n=10 mice per group, bars = mean, error bars = standard deviation. One-way ANOVA followed by post-hoc Tukey's test: *p<0.05, **p<0.01, ***p<0.001, n.s. = not significant.

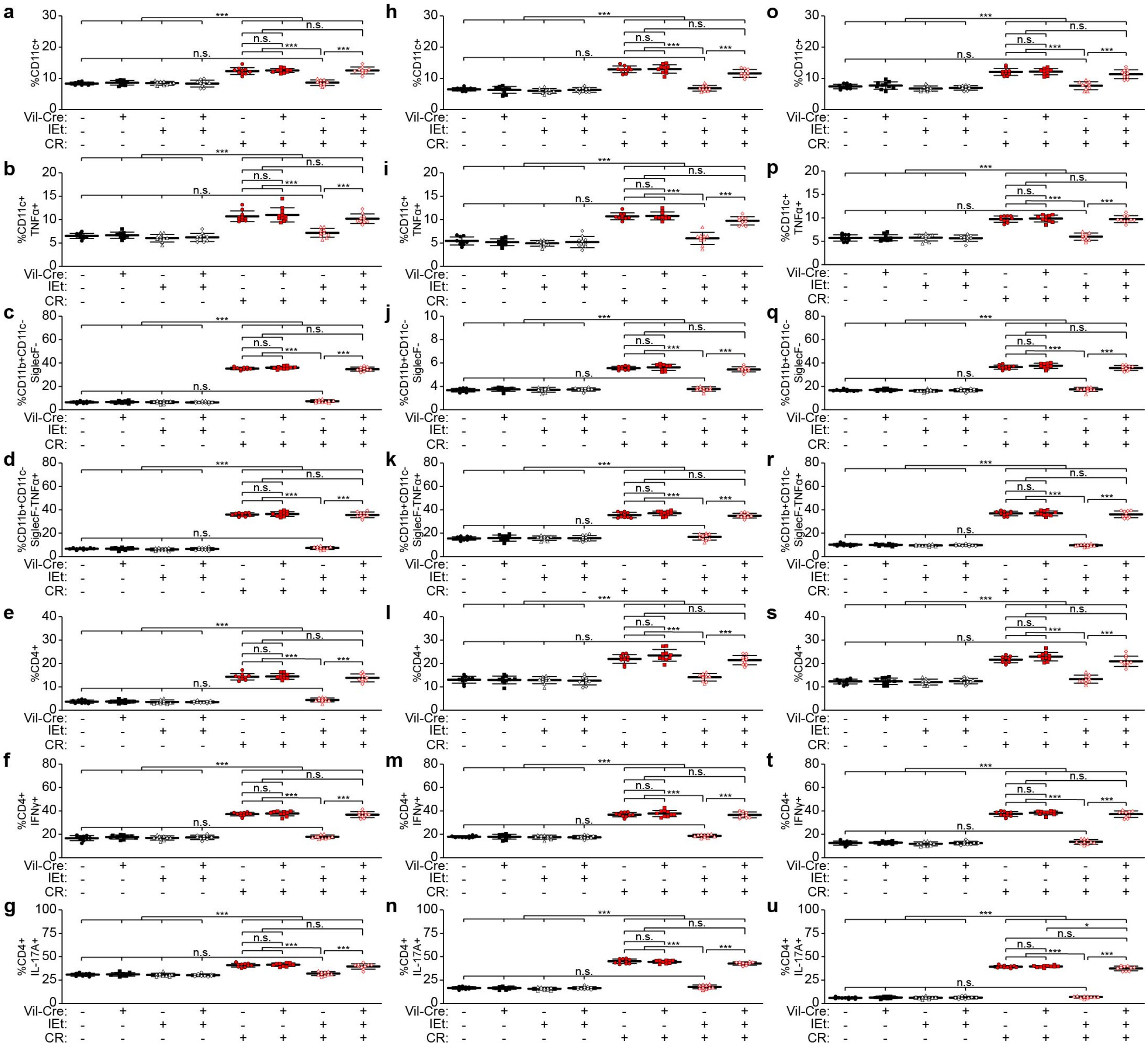


**Extended Data Fig. 13. IEt decreases immune cell infiltrates and inflammation via DRD2 expressed in IECs.** *Drd2*fl/fl x Villin (Vil)-Cre and *Drd2*fl/fl mice were administered IEt (600 mg/kg) daily by oral gavage for 2 d, and then infected with *C. rodentium* (CR, oral gavage, $10^8$ CFU) with continued metabolite treatment. Representative gating strategy, see Extended Data Fig. 10a. **a–u**, Indicated cell populations were analyzed from the (**a–g**) LP, (**h–n**) MLN, and (**o–u**) SP by FACS. Data are representative of at least 3 independent experiments, n=10 mice per group, bars = mean, error bars = standard deviation. One-way ANOVA followed by post-hoc Tukey's test: *p<0.05, ***p<0.001, n.s. = not significant.

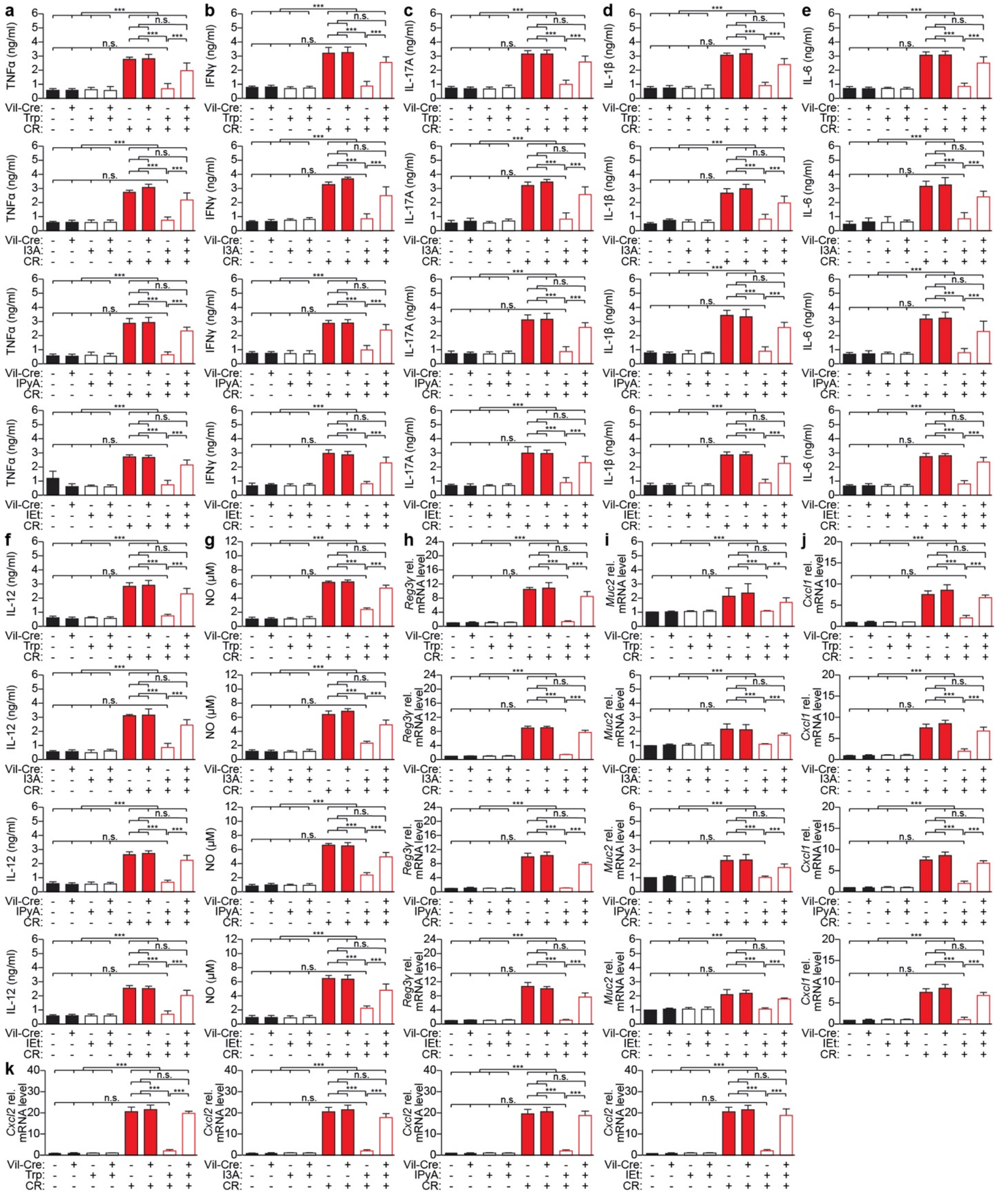
a
b
c
d
e
f
g
h
i
j
k
TNFα (ng/ml)
IFNγ (ng/ml)
IL-17A (ng/ml)
IL-1β (ng/ml)
IL-6 (ng/ml)
IL-12 (ng/ml)
NO (μM)
Reg3γ rel. mRNA level
Muc2 rel. mRNA level
Cxcl1 rel. mRNA level
Cxcl2 rel. mRNA level
Vil-Cre:
Trp:
I3A:
IPyA:
IEt:
CR:
n.s.
***
**

**Extended Data Fig. 14. Trp metabolites decrease inflammatory mediator production via DRD2 expressed in IECs.** *Drd2*fl/fl x Villin (Vil)-Cre and *Drd2*fl/fl mice were fed a conventional (2 g Trp/kg diet, ad libitum) or Trp (42 g Trp/kg diet, ad libitum) diet for 7 d or Trp metabolites (I3A (1000 mg/kg), IPyA (2900 mg/kg), or IEt (600 mg/kg)), by oral gavage daily for 2 d, and then infected with *C. rodentium* (CR, oral gavage, $10^8$ CFU) with continued Trp feeding or metabolite treatment. **a–g**, Conditioned media from the colon tissue was analyzed by (**a–f**) ELISA for the indicated cytokines. (**g**) Nitric oxide was measured by the Griess assay. **h–k**, Colon tissue cDNA was analyzed by qPCR for the indicated mRNA transcripts. Data are representative of at least 3 independent experiments, n=10 mice per group, bars = mean, error bars = standard deviation. One-way ANOVA followed by post-hoc Tukey's test: **p<0.01, ***p<0.001, n.s. = not significant.

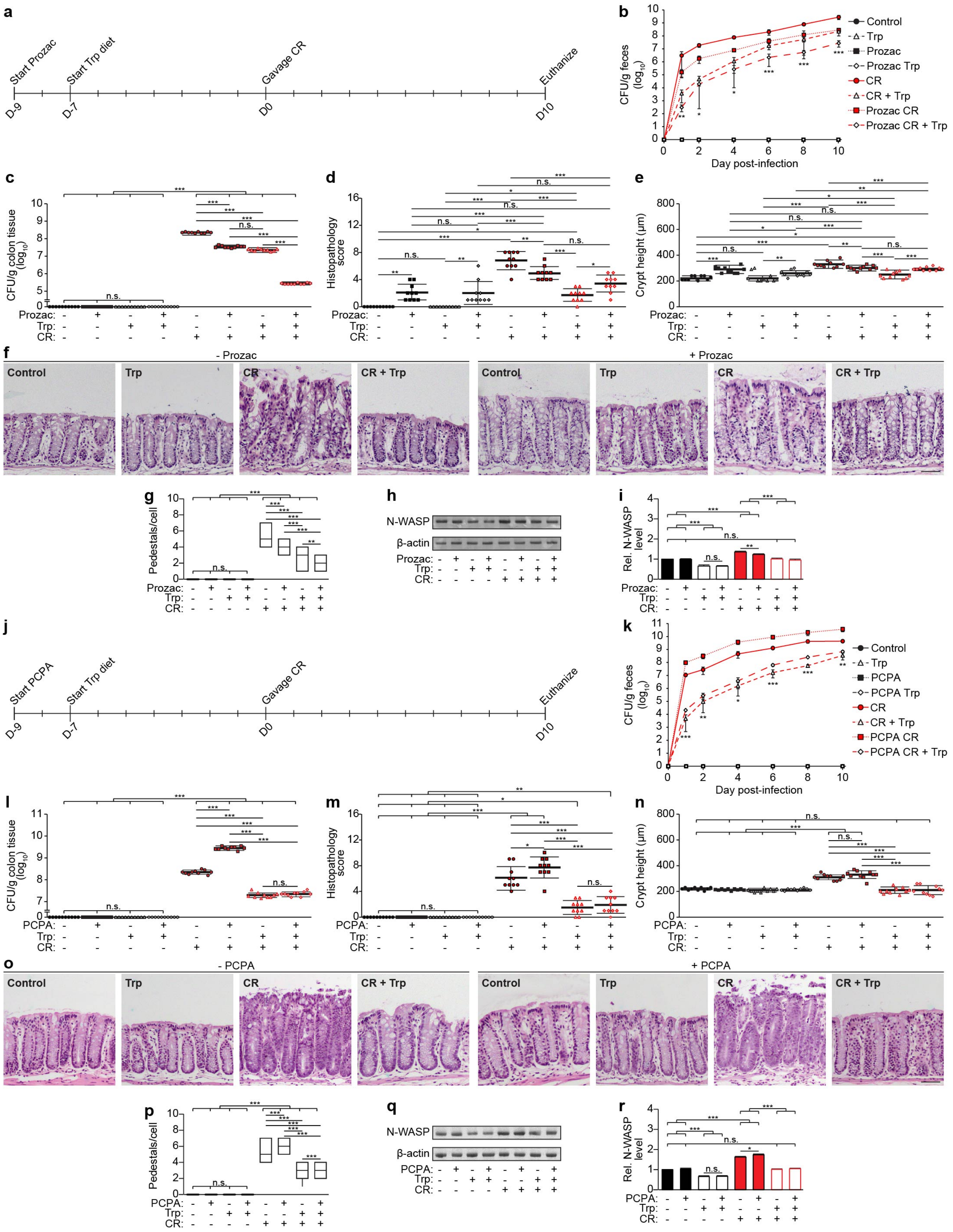

a
Start Prozac
Start Trp diet
Gavage CR
Euthanize
D-9
D-7
D0
D10
b
CFU/g feces (log10)
Day post-infection
Control
Trp
Prozac
Prozac Trp
CR
CR + Trp
Prozac CR
Prozac CR + Trp
c
CFU/g colon tissue (log10)
d
Histopathology score
e
Crypt height (μm)
Prozac:
Trp:
CR:
n.s.
f
- Prozac
+ Prozac
Control
Trp
CR
CR + Trp
g
Pedestals/cell
h
N-WASP
β-actin
i
Rel. N-WASP level
j
Start PCPA
Start Trp diet
Gavage CR
Euthanize
D-9
D-7
D0
D10
k
CFU/g feces (log10)
Day post-infection
Control
Trp
PCPA
PCPA Trp
CR
CR + Trp
PCPA CR
PCPA CR + Trp
l
CFU/g colon tissue (log10)
m
Histopathology score
n
Crypt height (μm)
PCPA:
Trp:
CR:
o
- PCPA
+ PCPA
Control
Trp
CR
CR + Trp
p
Pedestals/cell
q
N-WASP
β-actin
r
Rel. N-WASP level

**Extended Data Fig. 15. Trp diet provides protection again *C. rodentium* irrespective of serotonin levels.** C57Bl/6 mice were pre-treated with (**a–i**) serotonin selective reuptake transporter and tryptophan hydroxylase inhibitors, Prozac (20 mg/kg, oral gavage) and (**j–r**) 4-chloro-DL-phenylalanine methyl ester hydrochloride (PCPA, 150 mg/kg, oral gavage), to respectively increase or decrease serotonin levels, for 2 d, followed by conventional (2 g Trp/kg diet, ad libitum) or Trp (42 g Trp/kg diet, ad libitum) diet for 7 d. The mice were then administered *C. rodentium* (CR, oral gavage, $10^8$ colony-forming units, CFU) with continued Trp diet and inhibitor treatment. **b, c, k, l**, Bacterial load in (**b, k**) feces and (**c, l**) colon tissue was measured (**b, k**) every 1–2 d for 10 d post-infection or (**c, l**) at the peak of infection, 10 d post-infection. **d–f, m–o**, Colon sections were stained with H&E and (**d, m**) blindly scored for submucosal edema (0-3), goblet cell depletion (0-3), epithelial hyperplasia (0-3), epithelial integrity (0-4), and neutrophil and mononuclear cell infiltration (0-3). Data are expressed as the sum of these individual scores (0-16). See Methods for full description of scoring rubric. (**e, n**) Crypt heights were measured. (**f, o**) Representative images. Scale bar: 50 μm. (**g, p**) Pedestal formation = # of pedestals per host cell (Prozac, n = 127–178; PCPA, n = 126–156). **h, i, q, r**, Intestinal epithelial cells were isolated, and cell lysates were analyzed by Western blotting with the indicated antibodies. (**i, r**) Densitometry was performed using FIJI. Data are representative of at least 3 independent experiments, n=10 mice per group, bars = mean, error bars = standard deviation. One-way ANOVA followed by post-hoc Tukey's test: *p<0.05, **p<0.01, ***p<0.001, n.s. = not significant.

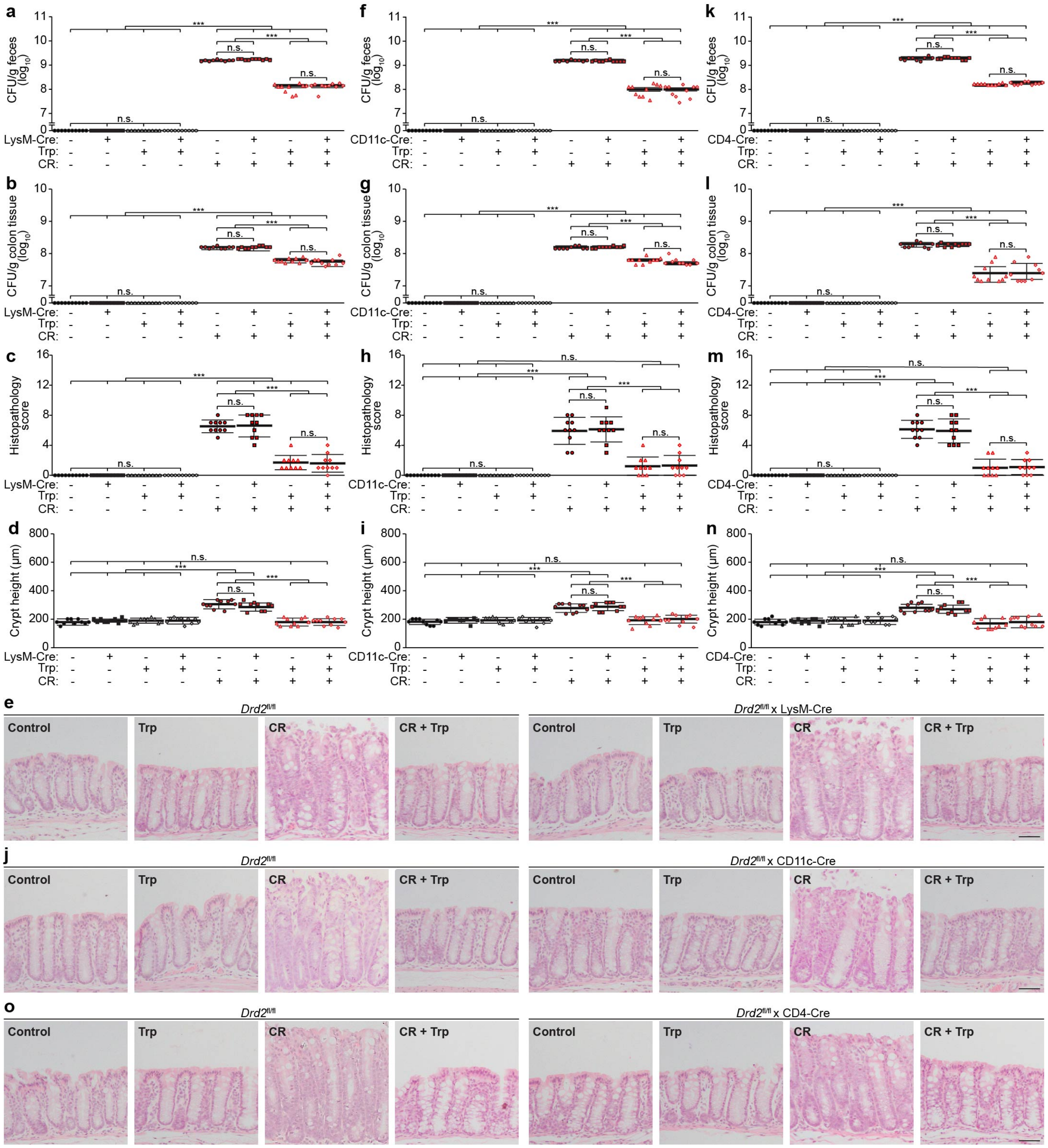

a
b
c
d
e
f
g
h
i
j
k
l
m
n
o
CFU/g feces (log10)
CFU/g colon tissue (log10)
Histopathology score
Crypt height (µm)
LysM-Cre:
CD11c-Cre:
CD4-Cre:
Trp:
CR:
n.s.
***
Drd2fl/fl
Drd2fl/fl x LysM-Cre
Drd2fl/fl x CD11c-Cre
Drd2fl/fl x CD4-Cre
Control
Trp
CR
CR + Trp

**Extended Data Fig. 16. Effects of the tryptophan (Trp) diet in protecting against *C. rodentium* infection do not depend on dopamine receptor D2 (DRD2) in macrophages, dendritic cells, or CD4$^+$ T cells. a–e,** *Drd2*fl/fl x LysM-Cre**, f–j**, *Drd2*fl/fl x CD11c-Cre, and **k–o,** *Drd2*fl/fl x CD4-Cre, and *Drd2*fl/fl mice were fed a conventional (2 g Trp/kg diet, ad libitum) or Trp (42 g Trp/kg diet, ad libitum) diet for 7 d and then infected with *C. rodentium* (CR, oral gavage, $10^8$ CFU) with continued Trp feeding. (**a, b, f, g, k, l**) Bacterial load in (**a, f, k**) feces and (**b, g, l**) colon tissue was measured at the peak of infection, 10 d post-infection. (**c–e**, **h–j, m–o)** Colon sections were stained with H&E and (**c, h, m**) blindly scored for submucosal edema (0-3), goblet cell depletion (0-3), epithelial hyperplasia (0-3), epithelial integrity (0-4), and neutrophil and mononuclear cell infiltration (0-3). Data are expressed as the sum of these individual scores (0-16). See Methods for full description of scoring rubric. (**d, i, n**) Crypt heights were measured. (**e, j, o**) Representative images. Scale bar: 50 μm. Data are representative of at least 3 independent experiments, n=10 mice per group, bars = mean, error bars = standard deviation. One-way ANOVA followed by post-hoc Tukey's test: ***$p<0.001$, n.s. = not significant.

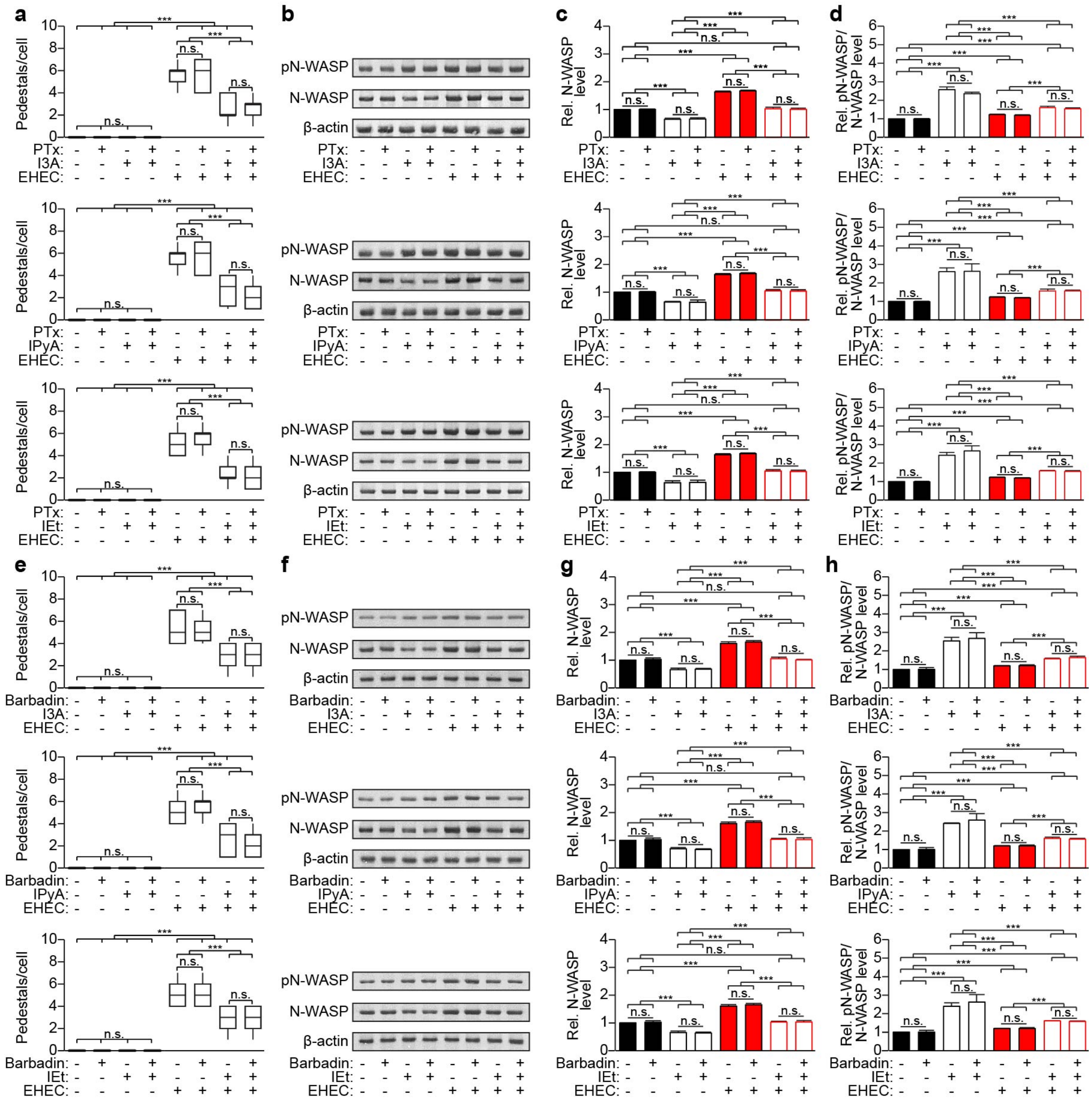

**Extended Data Fig. 17. Effects of Trp metabolites do not depend on Gαi and β-arrestin signaling.** Caco-2 cells were pre-treated with (**a–d**) Gαi inhibitor pertussis toxin (PTx, 100 ng/mL) for 18 h or (**e–h**) β-arrestin inhibitor barbadin (100 μM) for 30 min, followed by Trp metabolite (I3A, IPyA, or IEt, 100 μM) for 24 h. The cells were then infected with EHEC (MOI 50) for 12 h, (**a, e**) fixed, and stained with DAPI and Alexa Fluor 647-phalloidin. Pedestal formation = # of pedestals per Caco-2 cell (PTx: I3A, n=122–165; IPyA, n=113–146; IEt, n=105–141; barbadin: I3A, n = 125–141; IPyA, n = 126–166; IEt, n = 119–198). **b–d, f–h**, Alternatively, cells were lysed and analyzed by Western blotting with the indicated antibodies. (**c–d, g–h**) Densitometry was performed using FIJI. Data are representative of at least 3 independent experiments, n = 3, bars = mean, error bars = standard deviation. One-way ANOVA followed by post-hoc Tukey's test: ***p<0.001, n.s. = not significant.

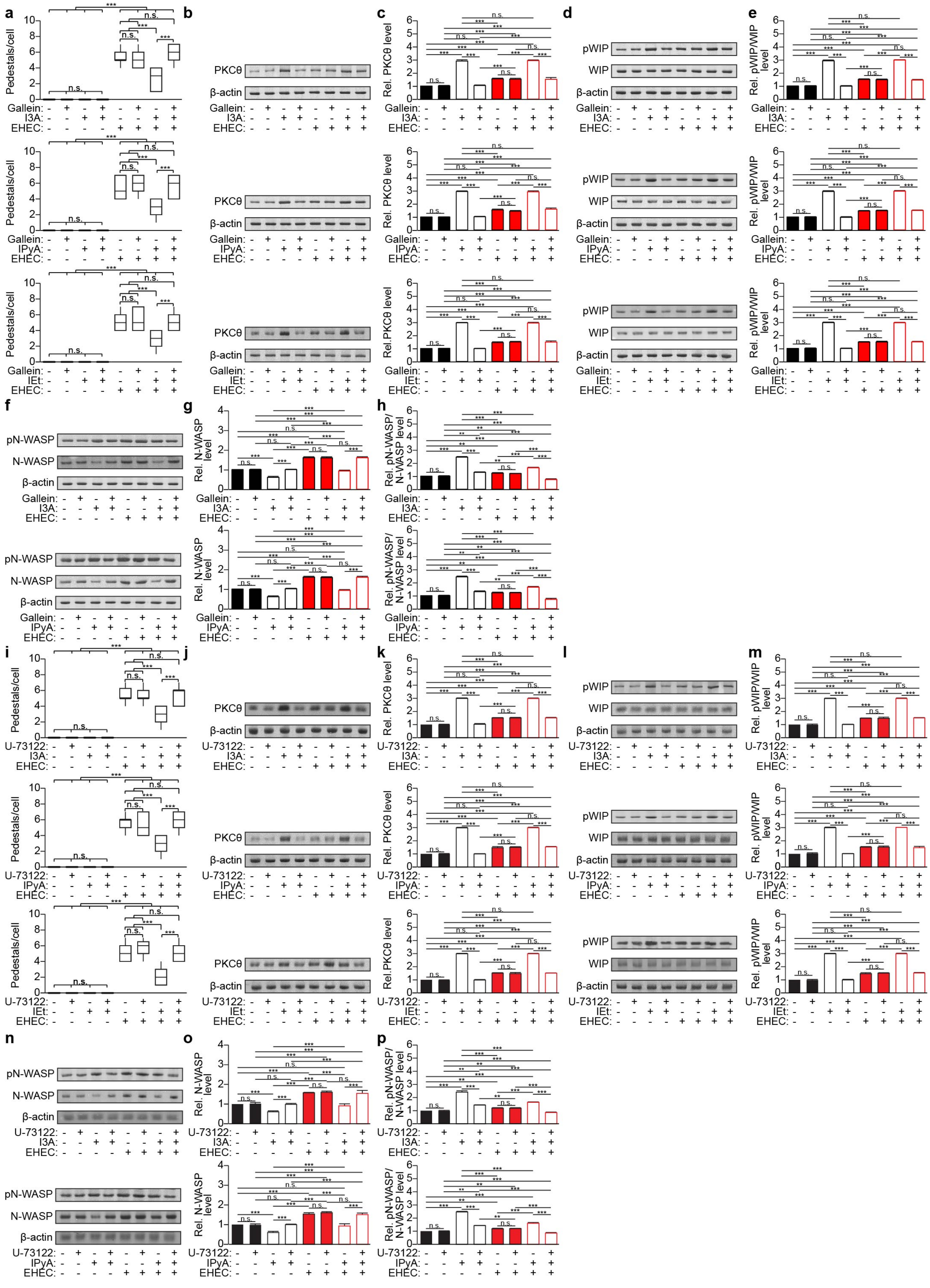
a
b
c
d
e
f
g
h
i
j
k
l
m
n
o
p
Pedestals/cell
PKCθ
β-actin
Rel. PKCθ level
pWIP
WIP
Rel. pWIP/WIP level
pN-WASP
N-WASP
Rel. N-WASP level
Rel. pN-WASP/N-WASP level
Gallein:
U-73122:
I3A:
IPyA:
IEt:
EHEC:
n.s.
***
**

**Extended Data Fig. 18. Effects of Trp metabolites depend on Gβγ and phospholipase C.** Caco-2 cells were pre-treated with (**a–h**) Gβγ inhibitor gallein (10 μM) for 30 min or (**i–p**) PLC inhibitor U-73122 (10 μM) for 30 min, followed by Trp metabolite (I3A, IPyA, or IEt, 100 μM) for 24 h. The cells were then infected with EHEC (MOI 50) for 12 h, (**a, i**) fixed, and stained with DAPI and Alexa Fluor 647-phalloidin. Pedestal formation = # of pedestals per Caco-2 cell (gallein: I3A, n=109–155; IPyA, n=157–185; IEt, n=118–133; U-73122: I3A, n=116–162; IPyA, n=143–182; IEt, n=128–195). **b–h, j–p**, Alternatively, cells were lysed and analyzed by Western blotting with the indicated antibodies. (**c, e, g, h, k, m, o, p**) Densitometry was performed using FIJI. Data are representative of at least 3 independent experiments, n = 3, bars = mean, error bars = standard deviation. One-way ANOVA followed by post-hoc Tukey's test: **p<0.01, ***p<0.001, n.s. = not significant.

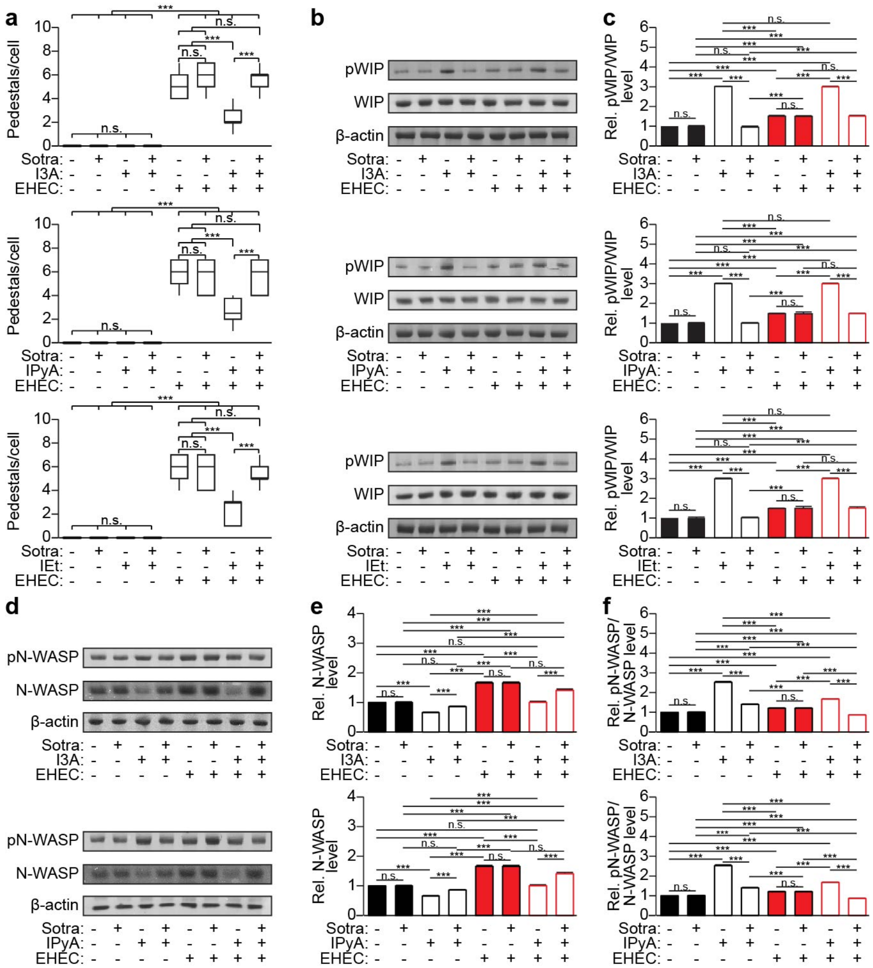

**Extended Data Fig. 19. Effects of Trp metabolites depend on protein kinase C (PKC).** Caco-2 cells were pre-treated with pan-PKC inhibitor sotrastaurin (Sotra, 5 μM) for 30 min, followed by Trp metabolite (I3A, IPyA, or IEt, 100 μM) for 24 h. The cells were then infected with EHEC (MOI 50) for 12 h, (**a**) fixed, and stained with DAPI and Alexa Fluor 647-phalloidin. Pedestal formation = # of pedestals per Caco-2 cell (I3A, n=123–134; IPyA, n=172–195; IEt, n=123–199). **b–f**, Alternatively, cells were lysed and analyzed by Western blotting with the indicated antibodies. (**c, e, f**) Densitometry was performed using FIJI. Data are representative of at least 3 independent experiments, n = 3, bars = mean, error bars = standard deviation. One-way ANOVA followed by post-hoc Tukey's test: ***p<0.001, n.s. = not significant.

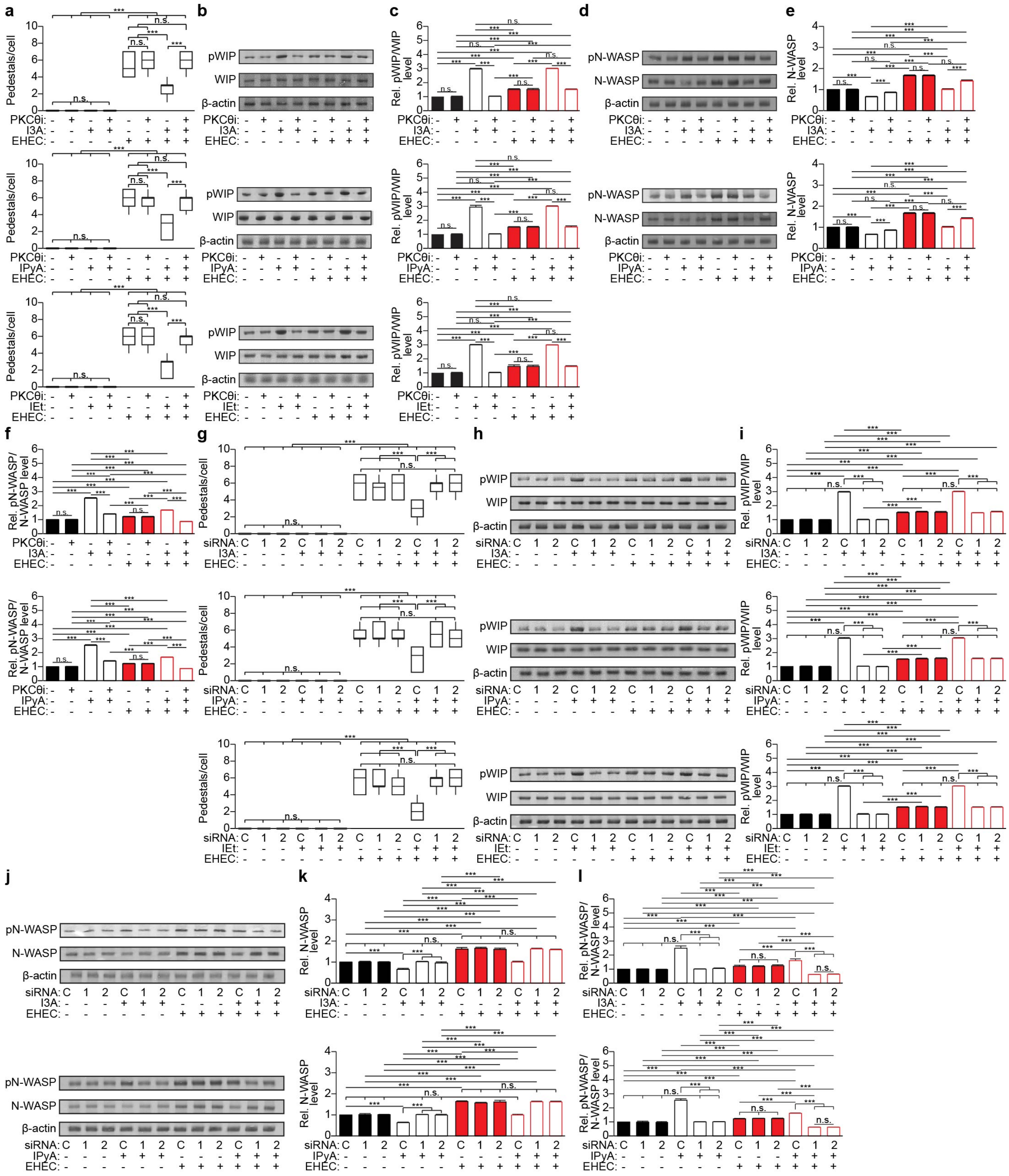
a
b
c
d
e
f
g
h
i
j
k
l
Pedestals/cell
pWIP
WIP
β-actin
Rel. pWIP/WIP level
pN-WASP
N-WASP
Rel. N-WASP level
Rel. pN-WASP/N-WASP level
PKCθi:
I3A:
IPyA:
IEt:
EHEC:
siRNA:
n.s.
***

**Extended Data Fig. 20. Effects of Trp metabolites depend on protein kinase C (PKC)-θ.** Caco-2 cells were pre-treated with (**a–f**) isoform-selective PKC-θ inhibitor (PKCθi, 5 μM) for 24 h, or (**g–l**) PKC-θ was knocked down using two different siRNA duplexes (1 and 2) or a negative control siRNA duplex (C), followed by Trp metabolite (I3A, IPyA, or IEt, 100 μM) for 24 h. The cells were then infected with EHEC (MOI 50) for 12 h, (**a, g**) fixed, and stained with DAPI and Alexa Fluor 647-phalloidin. Pedestal formation = # of pedestals per Caco-2 cell (PKCθi: I3A, n = 127–189; IPyA, n = 143–163; IEt, n = 157–160; siRNA: I3A, n = 170–196; IPyA, n = 118–159; IEt, n = 134–164). **b–f, h–l**, Alternatively, cells were lysed and analyzed by Western blotting with the indicated antibodies. (**c, e, f, i, k, l**) Densitometry was performed using FIJI. Data are representative of at least 3 independent experiments, n = 3, bars = mean, error bars = standard deviation. One-way ANOVA followed by post-hoc Tukey's test: ***p<0.001, n.s. = not significant.

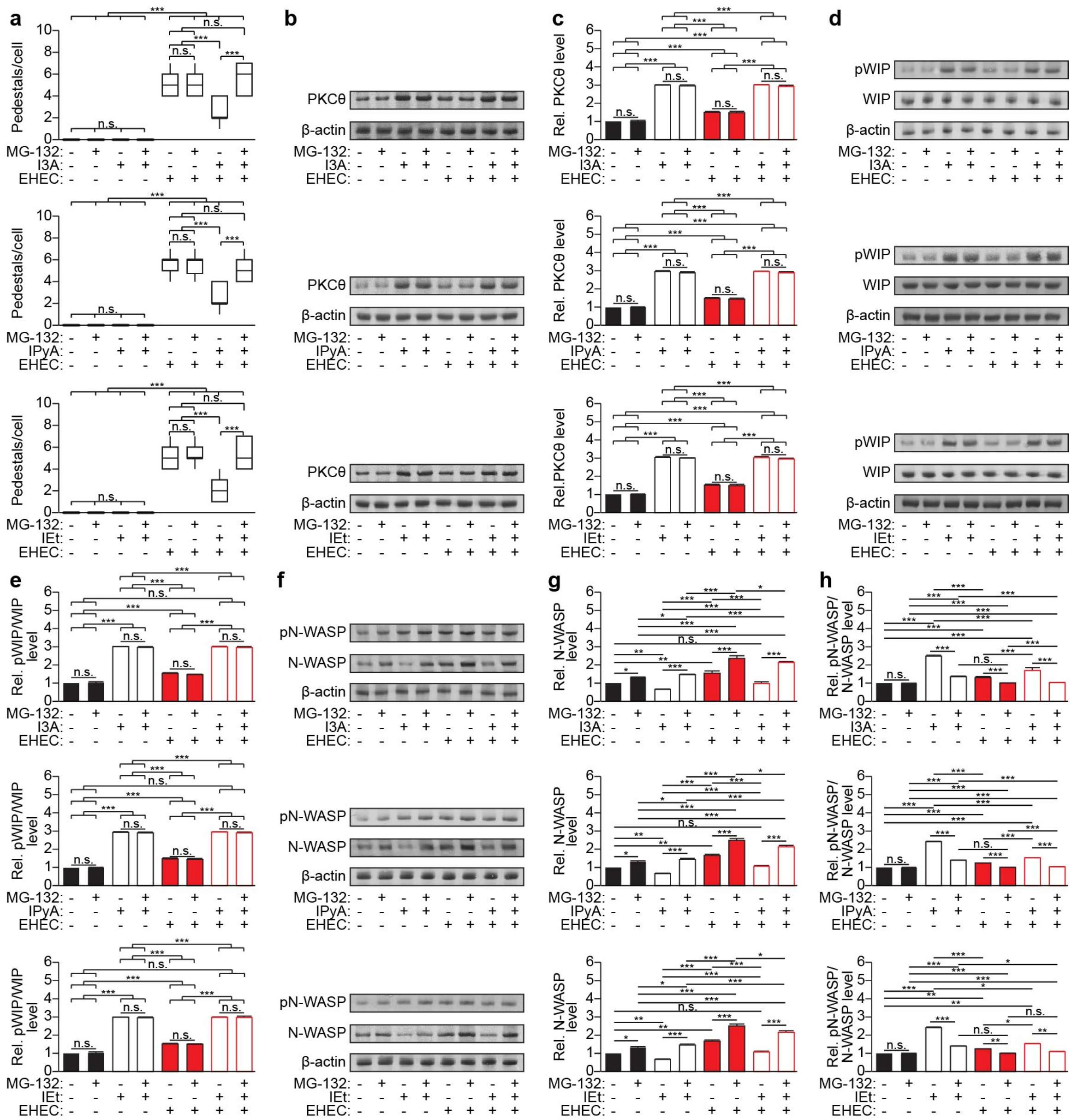

**Extended Data Fig. 21. Effects of Trp metabolites depend on proteasomal degradation.** Caco-2 cells were pre-treated with proteasomal inhibitor MG-132 (10 μM) for 1 h, followed by Trp metabolite (I3A, IPyA, or IEt, 100 μM) for 24 h. The cells were then infected with EHEC (MOI 50) for 12 h, (**a**) fixed, and stained with DAPI and Alexa Fluor 647-phalloidin. Pedestal formation = # of pedestals per Caco-2 cell (I3A, n=136–192; IPyA, n=129–168; IEt, n=113–137). **b–h**, Alternatively, cells were lysed and analyzed by Western blotting with the indicated antibodies. (**c, e, g, h**) Densitometry was performed using FIJI. Data are representative of at least 3 independent experiments, n = 3, bars = mean, error bars = standard deviation. One-way ANOVA followed by post-hoc Tukey's test: *p<0.05, **p<0.01, ***p<0.001, n.s. = not significant.
